# MYC pathway reprogramming through a TIP60 coactivator switch in neuroendocrine lineage transition in prostate cancer

**DOI:** 10.64898/2026.05.05.723058

**Authors:** Zhen Sun, Jimmy L. Zhao, Zahra F. Khan, Wazim M. Ismail, Teng Han, Subhiksha Nandakumar, Serina Young, Matthew Lange, Pan Cheng, Richard Koche, Nikolaus Schultz, Alexandre Gaspar-Maia, Charles L. Sawyers

## Abstract

Prostate adenocarcinomas (PRAD) can acquire resistance to androgen receptor signaling inhibitors through lineage transition to a cell state known as neuroendocrine prostate cancer (NEPC). Using a panel of isogenic PRAD and NEPC mouse tumoroids, we show that NEPC cells acquire new transcription factor (TF) dependencies that function in a previously undefined network. Through selective perturbation of each TF, we identify ASCL1 as a key regulator of NE lineage fate whereas MYCL functions downstream to drive NEPC growth/survival by recruitment of the TIP60/KAT5 acetyltransferase. Interestingly, while dependencies on specific TF family paralogs can vary across NEPC models, all show markedly enhanced dependency on TIP60. Moreover, the H2A.Z-acetyltransferase activity of the TIP60 complex (TIP60-C) is required for NEPC as well as the acetyl-reader BRD8, which is newly incorporated as a TIP60-C subunit with the NEPC transition. Targeted degradation studies in isogenic tumoroids reveal increased dependence on MYCL in NEPC relative to its paralog MYC in PRAD. In addition to a paralog switch (MYC to MYCL), the MYC pathway-addicted NE state is accompanied by a chaperone switch (from TIP60-C to SRCAP) for H2A.Z histone exchange and a coactivator switch (to TIP60) for MYC target gene expression. The NE-specific coupling of MYCL with TIP60 reveals a previously unappreciated opportunity to target MYC-driven NE diseases through pharmacological inhibition of TIP60.

## Introduction

Lineage plasticity is increasingly recognized as a mechanism of acquired resistance to targeted cancer therapies. PRAD and EGFR-mutant lung adenocarcinoma (LUAD) are compelling examples of an extreme form of lineage plasticity in which tumors transition from a canonical epithelial carcinoma to a neuroendocrine (NE) cancer under the selective pressure of targeted therapy, with androgen receptor signaling inhibitors (ARSI) and EGFR inhibitors respectively^1,2^. In both prostate and lung, lineage transition is a dynamic process, typically emerging in a genomic background of *RB1* and *TP53* loss, which together are thought to establish a cell state poised for lineage transition. In PRAD, the transition to NEPC requires induction of the master regulator NE TF ASCL1, accompanied by activation of additional TFs and changes in chromatin landscape^3–5^. Notably, the NE lineage state that emerges from PRAD shares remarkable similarity to that which emerges from EGFR-mutant LUAD, as well as *de novo* small cell lung cancer (SCLC), including expression of ASCL1, NEUROD1 and POU2F3, which define the major SCLC subgroups^6–8^. In all three NE disease settings, prognosis is poor despite extensive efforts to develop effective therapies.

The similarities across these NE diseases raises the intriguing possibility of common mechanism and dependencies. To date, much of the work exploring these questions has focused on SCLC, enabled by the availability of SCLC cell lines amenable for large scale, high throughput CRISPR screens. These efforts have yielded several druggable candidates including EZH2/EED, LSD1, KDM5A, KDM6A and others, several of which have progressed to clinical trials^9–12^. Of these, EZH2 inhibitors are the most clinically advanced but have shown limited efficacy in SCLC^13^. In contrast, such efforts in NEPC have been limited by a paucity of suitable cell line models, but recent work from our group and others suggests that prostate organoids are a useful platform for CRISPR screening^14–16^. In addition, comparative analysis of histone modifications in PRAD versus NEPC organoids led to the identification of NSD2 as a candidate NEPC target^17^. Furthermore, genetic evidence implicates additional TFs beyond ASCL1 as dependencies in human and mouse NEPC models, including FOXA1, FOXA2, ONECUT2, BRN2, PROX1 and NKX2.1^18–27^. Whether these TFs define distinct NE states/diseases or function in collaborative networks is poorly understood.

We recently developed a genetically defined mouse prostate organoid transplantation model that recapitulates the dynamics of the PRAD to NEPC transition in a way that accurately mirrors the human disease^3,28^. Here we leverage this *in vivo* platform to isolate and expand multiple isogenic lines of PRAD and NEPC tumoroids *ex vivo* at a scale amenable for extensive biochemical and functional studies. Through comprehensive epigenome profiling, we demonstrate dynamic enhancer reprogramming. We then use TF and coactivator-focused CRISPR screening, coupled with chemical genetic targeted protein degradation, to identify five key TF families required for NEPC lineage survival (ASCL1, FOXA, SOX, NFI, MYC). Short term knockout of each individual TF revealed a core network, with ASCL1 acting upstream to specify the NE lineage and MYCL functioning downstream to drive proliferation, partnering with TIP60-C/KAT5 for coactivation. Furthermore, we find that MYC undergoes a paralog swap (to MYCL or MYCN) and a coactivator switch (to TIP60-C) that results in a state of greatly enhanced MYCL/TIP60 dependency following the NEPC transition. These studies provide a potential therapeutic window for pharmacologic inhibition of TIP60/KAT5 in NEPC patients.

## Results

### Extensive enhancer reprogramming during PRAD to NEPC transition

To comprehensively characterize the chromatin state changes associated with the PRAD-to-NEPC transition, we built upon our earlier work using genetically modified primary prostate organoids to model lineage plasticity *in vivo*^3,28^, by deriving several isogenic pairs of PRAD and NEPC tumoroids. Specifically, we explanted tumors from mice transplanted with *Rb1^-/-^*, *Trp53^-/-^*, *Pten^-/-^* (hereafter called TKO) organoids, then expanded PRAD and NEPC TKO tumoroid lines through multiple passages at a scale amenable for biochemical characterization and CRISPR screening (see methods) (**Figures 1A and S1A**). Extensive histologic and molecular profiling of the isogenic organoid series confirmed stable expression of marker genes and signatures typical for the PRAD (e.g., ECAD, AR) and NEPC (e.g., ASCL1, SYP, DLL3) lineages in mouse models and patients^7,29–31^ (**Figures 1B-E and S1B-D).** In addition to confirming these lineage identities, transcriptomic analysis (RNA-seq) revealed activation of inflammatory JAK/STAT pathway genes in PRAD tumoroids, mirroring changes previously reported *in vivo* in adenocarcinomas in PtRP (TKO) mice prior to the NEPC transition^7^ (**Figure 1E**). Thus, these isogenic pairs provide a platform for extensive biochemical and functional characterization while maintaining the original lineage states seen *in vivo*.

**Figure 1.**
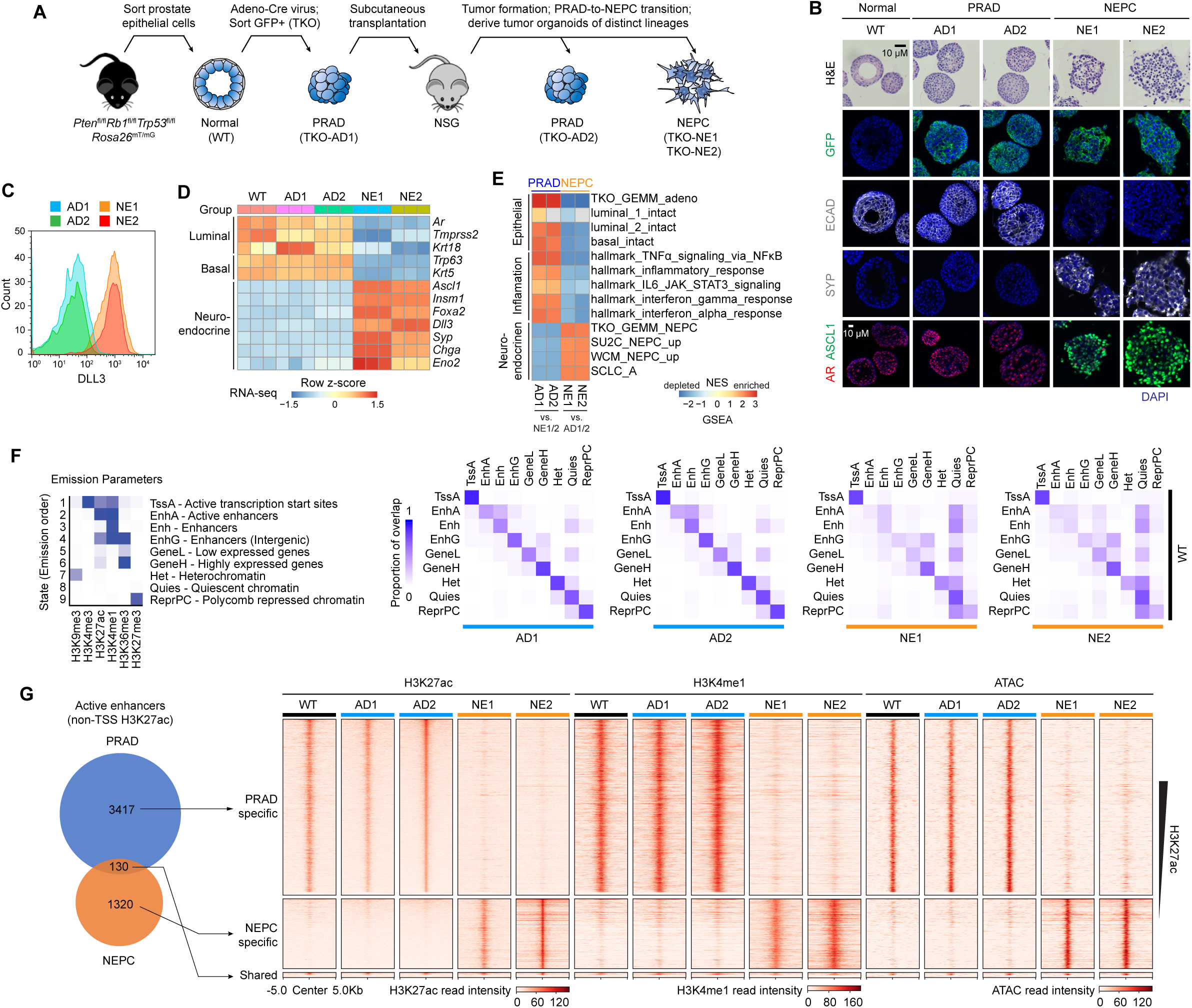
Extensive enhancer reprogramming during PRAD to NEPC transition. (**A**) Schematic showing the derivation of isogenic PRAD and NEPC tumor organoids following *in vivo* neuroendocrine transformation. (**B**) Hematoxylin and and eosin (H&E) and immunofluorescence (IF) of representative WT, PRAD and NEPC organoids. GFP indicates Cre-induced recombination (TKO). ECAD (E-cadherin), SYP (Synaptophysin). Scale bars representing 10 μm apply to all H&E and IF panels. (**C**) Flow cytometric analysis of DLL3 expression in PRAD and NEPC organoids. (**D**) Heatmap showing z-score transformed expression of luminal, basal and neuroendocrine marker genes from RNA-seq of indicated organoid lines. (**E**) Heatmap of normalized enrichment score (NES) from GSEA analysis of RNA-seq data showing significantly enriched gene sets in indicated PRAD and NEPC organoid lines (FDR < 0.05). Differentially expressed genes were identified by comparing individual PARD and NEPC line with both lines of the other lineage. Custom expression signatures of genes enriched in WT epithelial cells (luminal 1, luminal 2, basal) of intact mouse prostate^91^, genes enriched in PRAD or NEPC tumors from the TKO GEMM model^7^, genes upregulated in NEPC (vs. PRAD) patient tumor samples in two clinical cohorts (SU2C^31^ and WCM^29^), other neuroendocrine/neuronal signatures^7^, and hallmark inflammatory signaling signatures were analyzed. (**F**) (Left) Heatmap showing the emission parameters of the 9-state chromatin model built using chromoHMM with binarized CUT&RUN data from six histone modifications. The darker blue color indicates higher probability of observing the modification in the state. (Right) heatmap showing proportion of overlap (in bases) of chromatin states between tumor (PRAD and NEPC) and WT organoids across the genome as quantification of chromatin state changes. (**G**) (Left) Venn diagram showing overlap of non-TSS H3K27ac peaks, which are used to define active enhancers, in PRAD (AD2) and NEPC (NE2). (Right) heatmap showing enrichment of enhancer features (H3K27ac, H3K4me1, chromatin accessibility by ATAC-seq) at NEPC-specific, PRAD-specific and shared active enhancers across WT, PRAD and NEPC organoid lines.

Toward that end, we profiled six histone modifications (H3K4me1, H3K4me3, H3K36me3, H3K27ac, H3K9me3 and H3K27me3) commonly used to define the location and activity of different chromatin compartments genome-wide using CUT&RUN. ChromoHMM analysis revealed extensive epigenetic reprogramming following the NEPC transition, particularly at enhancer (active and poised) chromatin, whereas global promoter chromatin states remained largely similar (**Figures 1F**, **S1E, and S1F**). ATAC-seq, performed in parallel, revealed a profound redistribution of chromatin accessibility tracking that of enhancer modifications (H3K4me1 and H3K27ac) (**Figures 1G and S1G**).

### Network of transcription factor dependencies in murine and human NEPC

TFs such as ASCL1 are known to functionally reshape enhancer landscapes through pioneering activities^32,33^. To determine the relevance of ASCL1 and other TFs in our isogenic models, which may underlie the reprogrammed enhancer landscape, we used two criteria: (i) differential enrichment of TF binding motifs in accessible chromatin of NEPC versus PRAD tumoroids (**Figures 2A and S2A**) and/or (ii) differential expression in NEPC versus PRAD (**Figures 2B and S2B**). From this list of 25 candidates, we then performed arrayed CRISPR screening to assess the contribution of each TF to tumoroid fitness, scored by dropout of each TF-specific sgRNA (linked with blue fluorescent protein (BFP) expression) in a competition assay (**Figure 2C**). For each TF, 2-4 different sgRNAs were evaluated to ensure robustness of results (see methods). sgRNAs targeting the common essential genes *Mcm2* and *Pcna* served as positive controls, each showing <10% BPF expressing cells after 4 passages across two NEPC tumoroids (NE1, NE2) and PRAD tumoroids (AD2) (**Figure 2D**).

**Figure 2.**
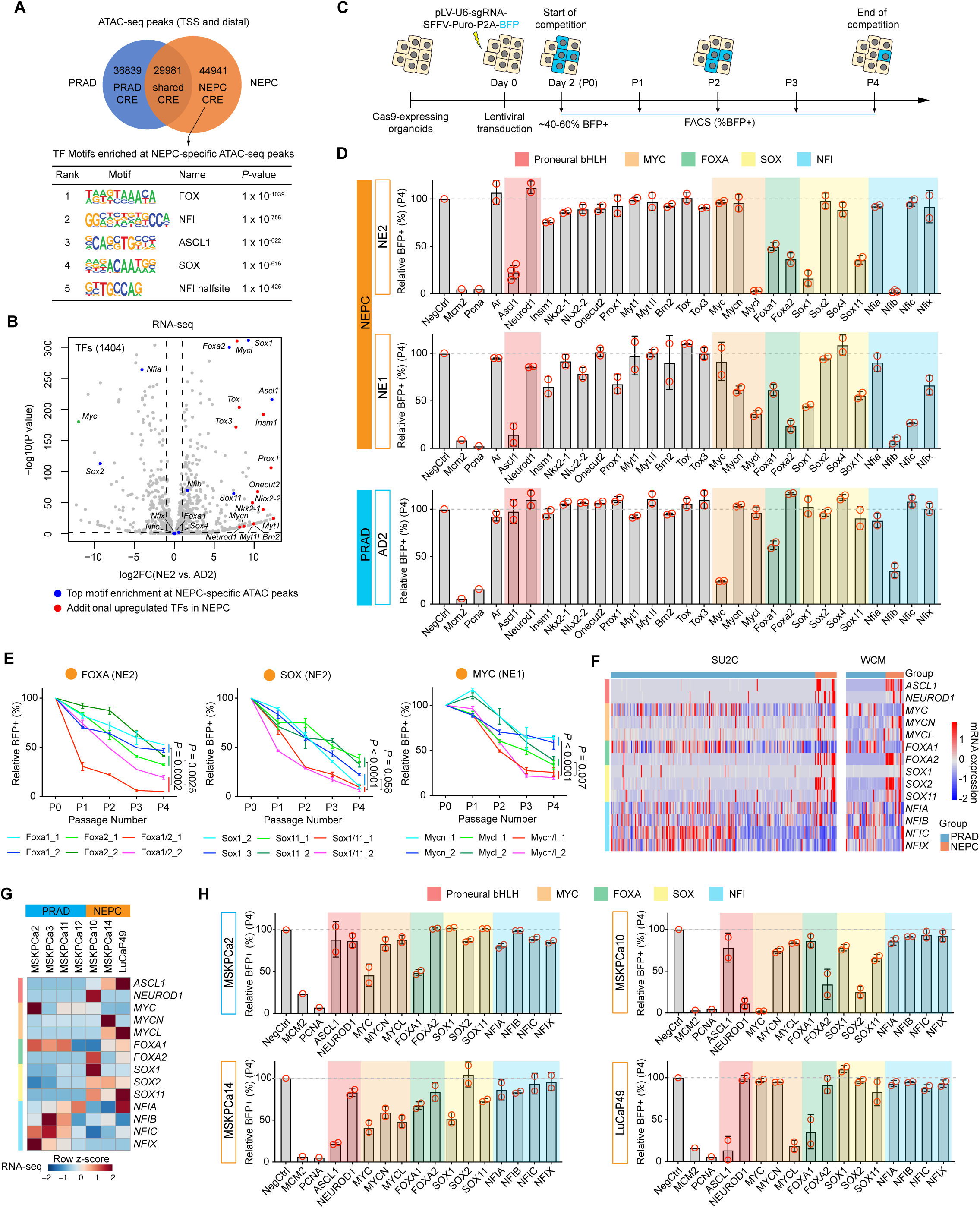
Network of transcription factor dependencies in murine and human NEPC. (**A**) (Top) Venn diagram showing overlap of ATAC-seq peaks in PRAD (AD1 and AD2) and NEPC (NE1 and NE2). CRE, *cis*-regulatory elements. (Bottom) Top 5 *de novo* TF motifs enriched at NEPC-specific ATAC-seq peaks (target coverage > 10%, ranked by *P* values). (**B**) Volcano plot showing differential RNA expression of TFs in NEPC (NE2) vs. PRAD (AD2). Screen candidates were highlighted in red and blue. (**C**) Schematic showing cell competition assay to assess cell proliferation upon knockout of individual genes. Percentage of BFP+ sgRNA-containing cells were monitored over a time course of 4 passages (P0-4). BFP percentage at day 2 (P0) was used as the baseline representing transduction efficiency. (**D**) Cell competition assays using at least two independent sgRNAs targeting the DNA binding domains of 25 individual TFs identified in (**A**) and (**B**) in indicated organoid lines, showing relative BFP percentages at P4 (day 18 for NE2 and AD2 and day 31 for NE1) after sgRNA transduction (see **Figure S2C** for measurements of individual sgRNAs and at all passages). BFP percentages were normalized to day 2 (P0) and the NegCtrl sgRNA at the respective day (relative %BFP+). The five families of NEPC lineage survival TFs were highlighted. sgRNAs targeting common essential genes implicated in DNA replication (*Mcm2* and *Pcna*) were used as positive controls. Dashed lines represent 100%. Each dot represents a sgRNA and is the average of two technical replicates. Data represent mean ± SD from n = 2 to 4 biological replicates (independent sgRNAs). (**E**) Cell competition assays using two independent single or dual sgRNA vectors targeting either or both TF paralogs in indicated NEPC tumoroids (see **Figure S2E** for PRAD), showing relative BFP percentage over 4 passages. BFP percentages were normalized to day 2 (P0) and the NegCtrl4 sgRNA at the respective day (relative %BFP+). Data represent mean ± SD from n = 2 technical replicates. *P* values were calculated using two-tailed unpaired Student’s *t*-tests. (**F**) mRNA expression (bulk RNA-seq) of key members of the five NEPC lineage survival TF families in histologically validated PRAD and NEPC tumor samples of two patient cohorts (SU2C and WCM). SU2C, n = 210 for PRAD and n = 22 for NEPC. WCM, n = 34 for PRAD and n = 15 for NEPC. (**G**) Heatmap showing mRNA expression (bulk RNA-seq) of key members of the five NEPC lineage survival TF families in indicated PDOs. (**H**) Cell competition assays using two independent sgRNAs targeting the DNA binding domains of key members of the five families of NEPC lineage survival TFs in indicated NEPC PDO lines, showing relative BFP percentages at P4 (day 31) after sgRNA transduction (see **Figure S2G** for measurements of individual sgRNAs and at all passages). BFP percentages were normalized to day 3 (P0) and the NegCtrl sgRNA at the respective day (relative %BFP+). sgRNAs targeting *MCM2* and *PCNA* were used as positive controls. Dashed lines represent 100%. Each dot represents a sgRNA and is the average of two technical replicates. Data represent mean ± SD from n = 2 biological replicates (independent sgRNAs).

As expected, we observed *Ascl1* dependency in both NEPC tumoroids, consistent with earlier reports in human and murine NEPC models^3,5^. Using <50% BFP-positive cells as a cutoff, six additional TFs (MYCL, FOXA1, FOXA2, SOX1, SOX11 and NFIB) also showed dependency in the NEPC tumoroids, with *Mycl*, *Nfib*, *Foxa2* and *Sox1* sgRNAs showing dropout levels comparable to (or greater than) *Ascl1* (**Figures 2D and S2C**). Of note, some TFs previously implicated in NEPC dependency (*Onecut2*, *Prox1, Nkx2-1, Insm1*)^21,22,24–26^ did not score in our tumoroid assay despite near complete protein knockout (**Figure S2D**), likely reflecting context-dependent heterogeneity across models. Five of the seven TF hits in our screen (*Ascl1*, *Mycl*, *Foxa2*, *Sox1*, and *Sox11*) are exclusively expressed in NEPC, whereas *Foxa1* and *Nfib* are expressed at comparable levels across both PRAD and NEPC lineages (**Figures 2B and S2B**). Indeed, we observe dependency on *Foxa1* and *Nfib* in both lineage states; however, the magnitude of dependency was higher in both NEPC tumoroids, likely a consequence of changes in the FOXA1 and NFIB cistromes after transitioning to the NE lineage (discussed later). Because both *Foxa1* and *Foxa2* are expressed in NEPC tumoroids and can function as paralogs, we asked if they compensate for each other in dependency assays by generating a dual sgRNA vector to simultaneously delete both genes. Indeed, NEPC tumoroids displayed significantly reduced survival after co-deletion of *Foxa1*/*Foxa2* compared to single gene deletion of *Foxa1* or *Foxa2* alone. We observed similar results following co-deletion of *Sox1* and *Sox11* compared to either of the single knockouts (**Figures 2E and S2E**).

In exploring the extreme *Mycl* dependency seen in NE2 tumoroids, we noted that the isogenic PRAD tumoroid (AD2) is dependent on the paralog *Myc* (*c-Myc*) whereas the NE1 tumoroids shows partial co-dependency on both *Mycl* and *Mycn*. Interestingly, these differences in paralog-specific dependency match the differential expression of each *Myc* paralog across the tumoroid panel (**Figure S2B**). Just as with *Foxa1/2* and *Sox1/11*, co-deletion of *Mycl* and *Mycn* in NE1 tumoroids revealed greater dependency than single knockout of either one alone (**Figures 2E and S2E**). Molecular details underlying this shift from *Myc* dependency in PRAD to *Mycl*/*Mycn* dependency in NEPC, including an unanticipated switch in coactivator usage, is discussed further below.

In sum, CRISPR screening of 25 candidate TFs across an isogenic panel of murine PRAD and NEPC tumoroid models identified NEPC-specific dependencies across five TF families (proneural bHLH, MYC, FOXA, SOX, NFI). To determine if these TF dependencies extend to human NEPC, we first confirmed expression of the relevant paralogs across two human castration resistant prostate cancer (CRPC) datasets enriched for NEPC (‘SU2C’^31^ and ‘WCM’^29^) (**Figures 2F and S2F**). The TF expression profiles largely mirror those seen in the murine tumoroids, except for SOX1 which is sporadically expressed in human NEPC (in contrast to SOX2). As expected, a subset of human NEPC is defined by expression of the proneural bHLH TF NEUROD1, largely mutually exclusive with ASCL1 as in human small cell lung cancer^6^.

To assess dependencies of these TFs in human NEPC, we performed CRISPR dropout screens in four patient derived organoids (PDOs)^7,34^ using the same BFP competition readout as in the murine tumoroids (**Figures 2G, 2H**, **and S2G**). The ASCL1+ (MSKPCa14, LuCaP49) and NEUROD1+ (MSKPCa10) models were selectively dependent on ASCL1 and NEUROD1 respectively, as expected, with the PRAD model MSKPCa2 serving as a control. We also observed dependency on MYC, FOXA and SOX family TFs within each model but with clear paralog-specific differences. For example, MYC family dependence in the ASCL1+ models mapped to MYCL (LuCaP49) or a combination of MYC paralogs including MYCL (MSKPCa14), whereas the NEUROD1 (MSKPCa10) and PRAD (MSKPCa2) models each showed exclusive MYC (c-MYC) dependency, largely consistent with expression levels of each paralog. Similar differences in paralog-specific dependency exist for FOXA and SOX family TFs. Of note, loss of any single NFI family TF was inconsequential for survival across all three NEPC models (and PRAD), perhaps due to compensation by other NFI paralogs co-expressed in human NEPC (in contrast to high *Nfib* expression in the mouse tumoroid model) (**Figure S2B**). While the findings in human NEPC models confirm many of the TF family dependencies identified in mice, they reveal considerable heterogeneity in the specific TF paralog dependencies within each family, perhaps due to heterogeneous expression and functional redundancy of multiple TF family paralogs.

### Enhanced dependency on the TIP60 coactivator complex in NEPC

Having established that the NEPC lineage transition is accompanied by a reshaped enhancer landscape coupled with a network of novel TF family dependencies, we next determined which coactivator enzymes were used by these TFs to induce gene expression. To address this question, we performed another arrayed CRISPR dropout screen, now targeting the catalytic domains of 25 different coactivator enzymes/complexes expressed in NEPC tumoroids **(Figure S3A)**, which together represent the chromatin regulatory pathways involved in ATP-dependent chromatin remodeling, DNA and histone methylation, and histone acetylation. For paralogous genes such as *Brg1*/*Brm*, *Ep300*/*Crebbp*, *Nsd1*/*Nsd2*, etc. where both paralogs were expressed (total of 8 pairs), we scored the consequences of single as well as double paralog knockout to address possible compensation. As with the TF screen, dependency was scored by progressive loss of BFP+ cells over 4 passages using 2-3 independent sgRNAs per target (**Figure 3A**). Common essential transcriptional regulator *Brd4* served as a dependency control in both PRAD and NEPC lineages, with each lineage showing similar degrees of dropout of BFP+ cells. As validation of the screening strategy, we noted selective dropout of *Brg1* sgRNAs and >800-fold increased sensitivity to the BRG1/BRM degrader AU-15330 in PRAD tumoroids compared to NEPC (**Figures 3A, 3B**, **and S3B**), consistent with prior evidence of BAF complex dependency in human CRPC (PRAD) models^35^.

**Figure 3.**
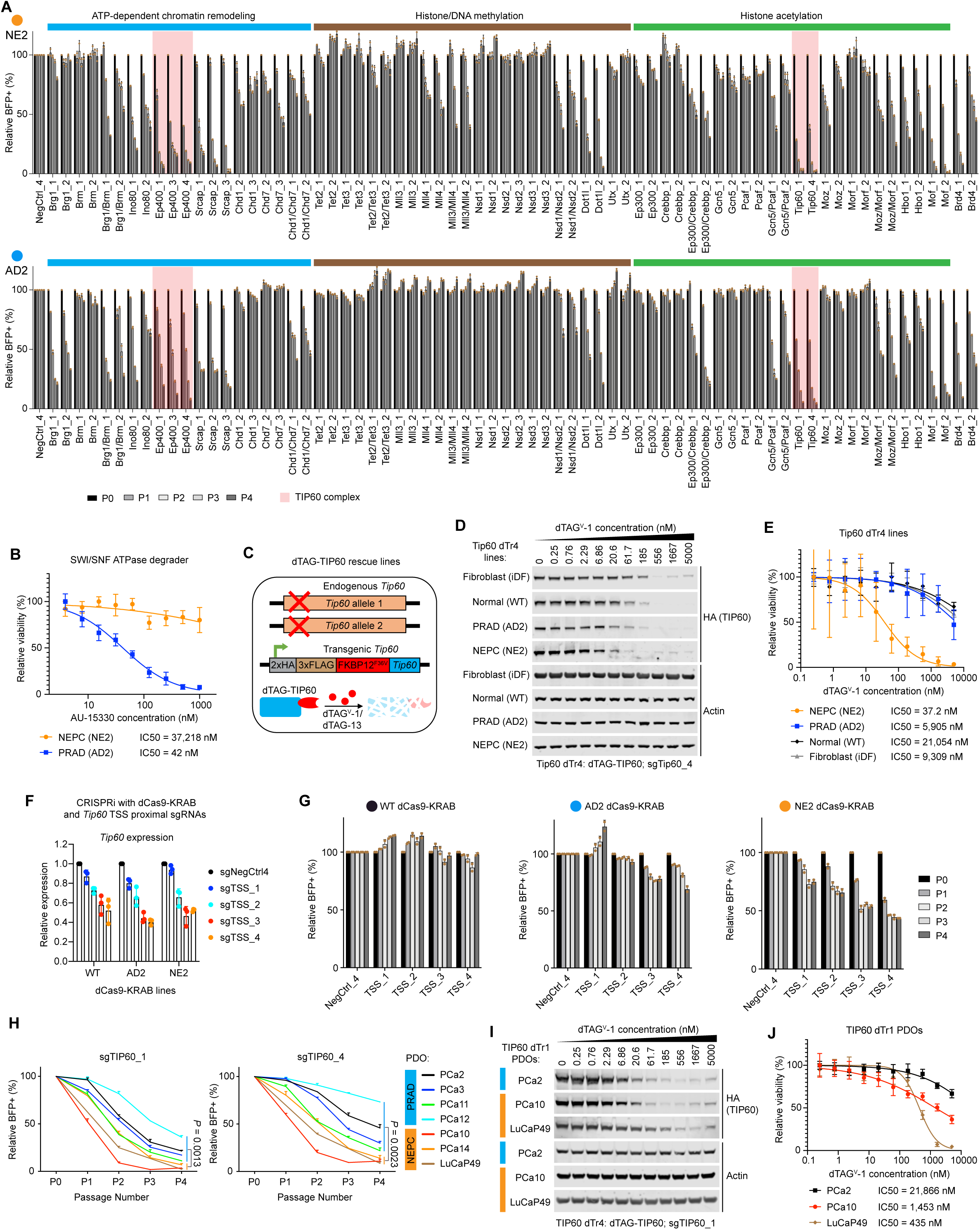
Enhanced dependency on the TIP60 coactivator complex in NEPC. (**A**) Cell competition assays using at least two independent single or dual sgRNA vectors targeting the catalytic domains of either or both paralogs of chromatin coactivator enzymes in indicated organoid lines, showing relative BFP percentage over 4 passages (P0-4) after sgRNA transduction. BFP percentages were normalized to P0 (day 2) and the NegCtrl4 sgRNA at the respective day (relative %BFP+). Subunits of the TIP60 complex (*Tip60* and *Ep400*) were highlighted (pink shade). sgRNAs targeting *Brd4* were used as positive controls. Data represent mean ± SD from n = 2 technical replicates. (**B**) BRG1/BRM degrader AU-15330 dose response curves and IC50 values for NEPC (NE2) vs. PRAD (AD2). Tumoroid lines were treated with the same AU-15330 gradient as in **Figure S3B** for 6 days before CellTiter Glo assays were performed to determine cell viability. Data represent mean ± SD from n = 10 biological replicates. (**C**) Schematic showing the strategy for chemical genetic targeted degradation of TIP60. dTAG-Tip60 lines were engineered by lentiviral expression of a dTAG-Tip60 transgene and knockout of endogenous *Tip60* alleles. Targeted proteolysis was induced by treatment of bifunctional degraders dTAG^V^-1 or dTAG-13 for proteasome dependent degradation. (**D**) Western blots of whole cell extract from indicated dTAG-Tip60 lines treated with a 3-fold gradient of dTAG^V^-1 for 6 hours. dTr4 indicates *Tip60* knockout with sgTip60_4 and rescue expression of dTAG-Tip60. Actin was used as loading control. (**E**) dTAG^V^-1 dose response curves and IC50 values for indicated dTAG-Tip60 lines (see **Figure S3F** for parental lines). Cell lines were treated with the same dTAG^V^-1 gradient as in (**D**) for 6 days before CellTiter Glo assays were performed to determine cell viability. Data represent mean ± SD from n = 10 biological replicates. (**F**) RT-qPCR analysis of *Tip60* RNA expression in indicated organoid lines stably expressing dCas9-KRAB and transduced with control or sgRNAs targeting the TSS proximal regions of the *Tip60* promoter. Data represent mean ± SD from n = 3 technical replicates. (**G**) Cell competition assays using four independent *Tip60* TSS-targeting sgRNAs in indicated organoid lines stably expressing dCas9-KRAB, showing relative BFP percentage over 4 passages (P0-4). BFP percentages were normalized to day 2 (P0) and the NegCtrl4 sgRNA at the respective day (relative %BFP+). Data represent mean ± SD from n = 2 technical replicates. (**H**) Cell competition assays using two independent *TIP60* sgRNAs in indicated PRAD and NEPC PDOs, showing relative BFP percentage over 4 passages (P0-4). BFP percentage was measured only on P0, P2 and P4 for LuCaP49. BFP percentages were normalized to day 3 (P0) and the NegCtrl4 sgRNA at the respective day (relative %BFP+). Data represent mean ± SD from n = 2 technical replicates. *P* values were calculated using two-tailed unpaired Student’s *t*-tests. (**I**) Western blots of whole cell extract from indicated dTAG-TIP60 lines treated with a 3-fold gradient of dTAG^V^-1 for 6 hours. dTr1 indicates *TIP60* knockout with sgTIP60_1 and rescue expression of dTAG-TIP60. Actin was used as loading control. (**J**) dTAG^V^-1 dose response curves and IC50 values for indicated dTAG-TIP60 PDOs (see **Figure S3I** for parental lines). PDO lines were treated with the same dTAG^V^-1 gradient as in (**I**) for 10 - 13 days before CellTiter Glo assays were performed to determine cell viability. Data represent mean ± SD from n = 10 biological replicates.

We reasoned that if the TF dependencies seen in NEPC are each mediated through distinct sets of coactivators, we should find numerous coactivator dependencies across the screen. Conversely, if the NEPC TF network functions through shared coactivation, we might see dependency on a single coactivator complex. Using a cutoff of <25% BFP+ cells after 4 passages, we identified four dependencies in NEPC: *Dot1l*, *Mof/Kat8*, *Srcap*, and the TIP60 complex (hereafter TIP60-C), containing the *Ep400* scaffold and the lysine acetyltransferase (KAT) subunit *Tip60*/*Kat5* (**Figure 3A**). Among these, dependence on *Tip60*/*Kat5* was particularly striking based on the speed of dropout (∼25% BFP+ cells at passage 1, eventually reaching <5% BPF+ cells by passage 4), with *Ep400* showing comparable depth and kinetics. The screen also revealed compensation by several chromatin modifying enzyme paralogs, with double knockout of *Brg1*/*Brm*, *Mll3*/*Mll4*, *Nsd1*/*Nsd2*, *Ep300*/*Crebbp*, *Gcn5*/*Pcaf*, *Moz*/*Morf* resulting in depletion to <50% BFP+ cells whereas the corresponding single gene knockouts had much more modest or no phenotypes (**Figure 3A**). In summary, although the screen revealed several chromatin modifying enzyme dependences, the TIP60/KAT5 complex emerged as the top candidate for deeper investigation due to the unexpected speed and magnitude of sgRNA dropout for both of the subunits (*Tip60*, *Ep400*) that were interrogated.

Toward that end, we noted that PRAD tumoroids also showed TIP60-C dependency of similar magnitude as NEPC tumoroids but with slower kinetics, reaching <25% BFP+ cells after 3 passages instead of just 1. This result is consistent with the fact that TIP60 is listed in DepMap as a common essential gene^36^. However, the differential dropout kinetics in NEPC versus PRAD tumoroids suggests that tumor cells may acquire enhanced dependency after transition to the NEPC lineage, raising the possibility of a therapeutic index with pharmacologic TIP60 inhibition. To address this question, we used a chemical genetic approach (dTAG)^37,38^ to selectively degrade TIP60 protein in NEPC versus PRAD tumoroids, as well as normal prostate organoids and dermal fibroblasts^39^ as normal tissue controls. After replacement of endogenous *Tip60* with the *dTAG-Tip60* allele (see methods), we documented dose dependent (dTAG^V^-1) degradation of TIP60 protein within 6 hours in all four models (**Figures 3C, 3D**, **and S3C-E**). Remarkably, NEPC tumoroids showed ∼150-fold increased sensitivity to TIP60 degradation in viability assays compared to PRAD tumoroids, with even greater differences relative to normal prostate organoids and dermal fibroblasts (**Figure 3E**). As expected, parental tumoroids lacking the *dTAG-Tip60* allele were insensitive to treatment with the dTAG^V^-1 degrader (**Figure S3F**). Consistently, TIP60 degradation leads to much more pronounced reduction in proliferation in NEPC tumoroids relative to other cell types in a cell competition-based assay (**Figure S3G**).

In exploring potential explanations for the large differential in IC50, we noted substantial TIP60 protein degradation in all four models at degrader concentrations flanking the proliferation IC50 in NEPC tumoroids (∼20-60nM), yet the other three cell lines showed no proliferation defect. This suggests that NEPC cells require a higher threshold of TIP60 protein than the other cell types we evaluated. To investigate this possibility, we used four different sgRNAs targeting the *Tip60* transcription start site (TSS), coupled with CRISPRi, to dampen the transcriptional output of the endogenous *Tip60* alleles (peaking at ∼50% reduction) (**Figure 3F**). As with the *dTAG*-*Tip60* allele, proliferation of NEPC tumoroids was impaired by modest reductions in *Tip60* mRNA levels whereas PRAD and normal prostate organoids were unaffected (**Figure 3G**).

Given the dramatic difference in TIP60 dependency observed in the murine PRAD versus NEPC models, we next asked if this enhanced TIP60 dependency extends to human NEPC. To do so, we measured the consequences of *Tip60* knockout across the 3 NEPC PDOs used earlier for TF CRISPR screening but now including an additional 4 PRAD PDOs to compare relative *Tip60* dependency. As with the murine tumoroids, *Tip60* sgRNAs were depleted across both lineages, but the kinetics of depletion was consistently faster in the NEPC lines, reaching <50% BFP+ cells by passage 1 or 2, compared to passage 3 or 4 for the PRAD lines (**Figures 3H and S3H**). As further confirmation of differential sensitivity to *Tip60* depletion, we engineered the *dTAG*-*Tip60* allele into representative PRAD (MSKPCa2) and NEPC (LuCaP49 and MSKPCa10) PDOs and again documented a dose-dependent decrease in TIP60 protein levels in both lineages (**Figure 3I**). As with the murine tumoroid models, both ASCL1+ (LuCaP49) and NEUROD1+ (MSKPCa10) NEPC PDOs showed enhanced sensitivity to TIP60 degradation in viability assays (∼50 and 15-fold, respectively) compared to the PRAD PDO (**Figures 3J and S3I**).

To summarize, CRISPR screening of 25 chromatin modifying enzymes expressed in the NEPC lineage state revealed 4 major dependencies, of which *Tip60* and the TIP60-C scaffold *Ep400* displayed the most profound dropout phenotype. Using chemical genetic targeted protein degradation and CRISPRi, we find that the proliferation of mouse and human NEPC cells is selectively impaired by modest (∼50%) reductions in TIP60 protein (dTAG) or mRNA levels (CRISPRi) relative to PRAD cells, suggesting the NEPC lineage has an enhanced need for TIP60-C activity compared to other cell lineages.

### TIP60-C dependency in NEPC is mediated through H2A.Z acetylation and selective BRD8 incorporation

Having established that NEPC requires a critical threshold level of TIP60 for survival, we next asked what TIP60 associated activities are responsible for this dependency. TIP60 is one component of a >1.5 megadalton complex consisting of ∼17 subunits^40–43^ (**Figures 4A and S4A**). TIP60-C is a potent transcriptional coactivator^44–48^ and has two major chromatin-associated enzymatic activities: (i) acetylation of histones H4 and H2A.Z (by the TIP60 KAT domain) and (ii) exchange of the histone variant H2A.Z in place of the canonical histone H2A (by EP400, an ATPase domain containing scaffold protein)^40–43,49^. TIP60-C can potentially also recognize chromatin states through protein subunits such as BRD8 and YEATS4 that contain histone reader domains. Of note, TIP60-C shares subunits with a second chromatin remodeler SRCAP that also promotes H2A-H2A.Z histone exchange but does not contain a histone acetylation module^43,50^ (**Figure 4A and S4A**).

**Figure 4.**
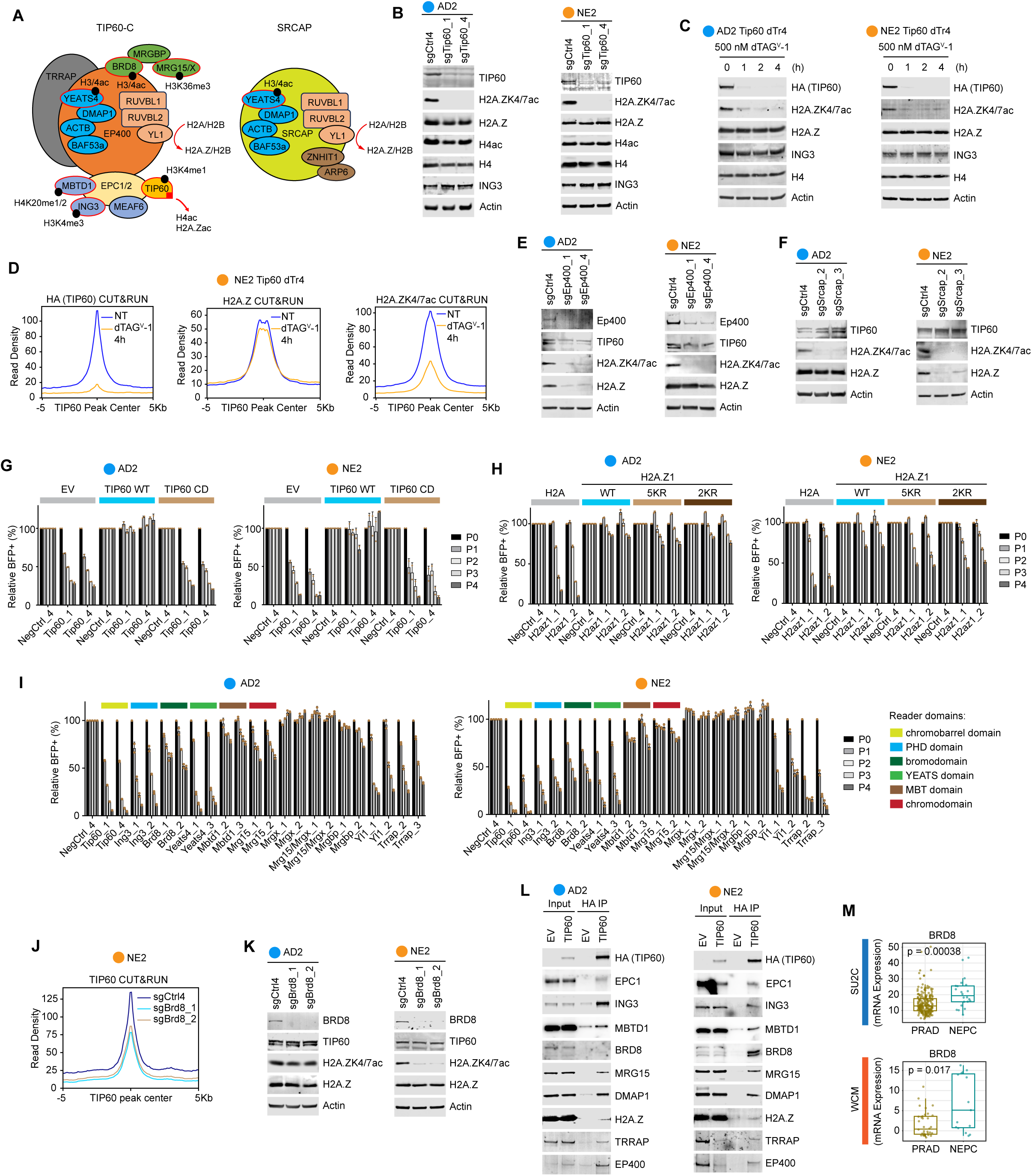
TIP60-C dependency in NEPC is mediated through H2A.Z acetylation. (**A**) Schematic showing the TIP60 and related SRCAP complexes. Red square indicates acetyltransferase domain. Red outlines indicate subunits with chromatin reader domain(s). Black circles indicate histone modifications reportedly/typically recognized by respective reader subunits. (**B**) Western blots of whole cell extract from AD2 and NE2 organoids with Tip60 knockout. In (**B**), (**C**), (**E**), (**F**), and (**K**), Actin was used as loading control. (**C**) Western blots of whole cell extract from NE2 dTAG-Tip60 (dTr4) organoids treated with 500 nM dTAG^V^-1 for indicated time. (**D**) Average profile showing occupancy (CUT&RUN) of TIP60 (via HA-tag), H2A.Z and H2A.Zac at TIP60 peaks in NE2 dTAG-Tip60 (dTr4) organoids treated with 500 nM dTAG^V^-1 for 4 h. NT, non-treated. (**E**) Western blots of whole cell extract from AD2 and NE2 organoids with *Ep400* knockout. (**F**) Western blots of whole cell extract from AD2 and NE2 organoids with *Srcap* knockout. (**G**) Cell competition assays using two independent *Tip60* sgRNAs in parental AD2 and NE2 tumoroids or those overexpressing wild type (WT) or Q377E/G380E catalytically dead (CD) TIP60. In (**G**), (**H**), and (**I**), data represent relative BFP percentage over 4 passages (P0-4). BFP percentages were normalized to day 2 (P0) and the NegCtrl4 sgRNA at the respective day (relative %BFP+). Data represent mean ± SD from n = 2 technical replicates. (**H**) Cell competition assays using two independent H2az1 sgRNAs in AD2 and NE2 organoids overexpressing an H2A/H2A.Z series. 2KR, K4/7R mutant. 5KR, K4/7/11/13/15R mutant. (**I**) Cell competition assays using two independent single or dual sgRNA vectors targeting individual TIP60 complex subunits or both paralogs in AD2 and NE2 organoids. Reader domain targeting sgRNAs were utilized for *Ing3*, *Brd8*, *Yeats4*, *Mbtd1*, and *Mrg15*. (**J**) Average profile showing occupancy (CUT&RUN) of TIP60 in NE2 organoids with *Brd8* knockout. NT, non-treated. (**K**) Western blots of whole cell extract from AD2 and NE2 organoids with *Brd8* knockout. (**L**) Western blot analyses of immunoprecipitation with HA tag in AD2 and NE2 organoids expressing HA-FLAG-tagged Tip60 (Tip60 dTr4) or empty vector (EV). (**M**) Boxplots showing mRNA expression (bulk RNA-seq) of *BRD8* in tumor samples of the SU2C and WCM prostate cancer patient cohorts. *P* values were calculated using two-tailed Wilcoxon rank sum test.

To assess the role of these three different TIP60-C activities, we first looked for changes in H4 and H2A.Z acetylation following *Tip60* knockout (by CRISPR) or TIP60 degradation (by dTAG) in PRAD and NEPC tumoroids. H2A.Z acetylation (H2A.Zac) was rapidly and almost completely abrogated upon loss of TIP60 across lineages, whereas H4 acetylation remained largely unaltered (**Figures 4B, 4C**, **and S4B**). CUT&RUN experiments confirmed colocalization of TIP60 and H2A.Z acetylation at enhancers and promoters in NEPC tumoroids (**Figure S4C**) and loss of H2A.Z acetylation within 4 hours of TIP60 degradation without affecting H2A.Z chromatin incorporation (**Figure 4D**), thereby defining TIP60 as the major H2A.Z acetyltransferase in NE cells.

To determine whether TIP60-C also mediates H2A-H2A.Z exchange activity in our models, we measured H2A.Z levels in PRAD and NEPC tumoroids after *Ep400* knockout. Interestingly, we observed near complete loss of H2A.Z protein in PRAD tumoroids but not in NEPC tumoroids (**Figure 4E**). Conversely, knockout of the analogous ATPase domain containing subunit in SRCAP resulted in loss of H2A.Z exchange in NEPC tumoroids but had no effect in PRAD tumoroids (**Figure 4F**). Thus, TIP60-C is responsible for H2A-H2AZ exchange activity in the PRAD lineage whereas SRCAP performs this function after the transition to NEPC.

Having shown that TIP60 is the critical acetyltransferase responsible for H2A.Z acetylation, we next asked if this activity is required for NEPC survival using two approaches. First, we engineered expression of a wildtype (WT) or catalytically dead allele of *Tip60* (Q377E/G380E)^51^ in NEPC tumoroids (**Figure S4D**) then performed rescue experiments by assessing CRISPR dropout of sgRNAs targeting the endogenous *Tip60* gene (linked with BFP, as in the previous dropout assays). Dropout of *Tip60* sgRNA was rescued by expression of catalytically active *Tip60* in both PRAD and NEPC cells, as expected, but not by the catalytically dead allele (**Figure 4G**), demonstrating TIP60 acetyltransferase activity is essential for survival in both lineages.

For the second approach, we asked if the H2A.Z acetylation mark itself, which associates with the gene activating role of H2A.Z^50,52,53^, is required for NEPC survival by replacement of endogenous H2A.Z with mutants that cannot be acetylated due to lysine-to-arginine (KR) substitutions at key substrate residues^50,54^. Because H2A.Z is encoded by two genes in the mammalian genome (*H2az1*, *H2az2*), we first performed selective CRISPR deletion of each to determine their relative importance for NEPC survival. sgRNA dropout assays revealed dependency on *H2az1*, but not *H2az2*, in both PRAD and NEPC (**Figure S4E**). Having established *H2az1* as the functionally relevant paralog, we engineered KR substitutions at K4/7 (2KR) and at K4/7/11/13/15 (5KR) and then performed cDNA rescue experiments following *H2az1* knockout, analogous to those performed with the catalytically dead *Tip60* allele. The 2KR and 5KR mutants showed loss of H2A.Z acetylation in PRAD and NEPC cells in both lineages (**Figure S4F**). However, in the *H2az1* sgRNA dropout assay, the 2KR and 5KR substitutions fully restored proliferation in PRAD tumoroids but showed only modest restoration in NEPC cells (**Figure 4H**). In summary, TIP60 acetyltransferase activity is required for survival in both PRAD and NEPC lineages but H2A.Z is a critical functional substrate only in the NEPC lineage.

In addition to its acetyltransferase and H2A.Z exchange activities, TIP60-C has the potential to bind chromatin directly through subunits such as BRD8 and YEATS4 that contain reader domains that recognize acetylated histone residues. To determine whether this chromatin binding activity of TIP60-C is important in NEPC, we repeated the *Tip60* CRISPR dropout assay but now measuring dependency on each of the reader domain containing subunits (BRD8, YEATS4, ING3, MBTD1, MRG15) as well as several scaffolding/adapter subunits (MRGX, MRGBP, YL1, TRAPP) (**Figures 4A and S4A**). NEPC and PRAD proliferation was severely impaired by knockout of *Brd8*, *Yeats4* and *Ing3*, whereas *Mbtd1* and *Mrg15* were largely dispensable (**Figure 4I**). Interestingly, the three essential reader subunits all contain domains that typically recognize histone modifications associated with active promoters/enhancers (H3/4ac, H3K4me3)^55–57^, whereas those of the non-essential subunits recognize DNA damage (H4K20me1/2) and gene body (H3K36me3) associated marks^58,59^. *Yl1* and *Trapp* also scored as dependencies, consistent with their adapter functions to interact with H2A.Z and TFs, respectively. TRRAP is of particular interest due to its ability to bind TFs such as MYC^60,61^ (discussed below).

Because the BRD8 (Bromo) and YEATS4 (YEATS) reader domains recognize histone acetylation marks^55,57^, which are typically present at both enhancers and promoters similarly as TIP60 (**Figure S4C**), we asked if either or both directly contribute to TIP60 chromatin binding. Indeed, CRISPR knockout of either *Brd8* or *Yeats4* resulted in a measurable decline in TIP60 occupancy in NEPC tumoroids, albeit not as pronounced as with *Tip60* degradation (**Figures 4J**, **S4G, and S4H**). Commensurate with reduced TIP60 binding, *Brd8* and *Yeats4* knockout also resulted in a pronounced decrease in H2A.Zac (**Figures 4K and S4I**). Interestingly, the reduction in H2A.Zac observed following *Brd8* knockout was restricted to NEPC tumoroids. To explore why *Brd8* depletion selectively impairs H2A.Zac in the NEPC lineage, we compared the subunit composition of TIP60-C in PRAD versus NEPC through side-by-side coimmunoprecipitation experiments. To our surprise, we found that BRD8 was present in TIP60-C in NEPC but not in PRAD (**Figure 4L**) and not in normal prostate organoids (**Figure S4J**), whereas all other TIP60-C subunits examined were equally present. Interestingly, in two independent patient cohorts, *BRD8* mRNA expression is higher in NEPC than PRAD (**Figures 4M and S4K)**. Of note, prior work defining the components of mammalian TIP60-C has been focused on model cell lines typically used for larger scale biochemical purification (K562^40,42,43,58,62^, HeLa^51,63,64^, and HEK293T^41^). To our knowledge, this is the first evidence that TIP60-C core subunit composition may vary across lineages.

Taken together, NEPC is dependent on the acetyltransferase activity (TIP60) and chromatin binding activity of TIP60-C (particularly via NE-enriched subunit BRD8), all of which are required for H2A.Z acetylation. Furthermore, mutagenesis studies implicate H2A.Z as a critical TIP60-C substrate required for NEPC survival. Interestingly, the H2A-H2A.Z exchange activity of TIP60-C, while present in PRAD, is lost during the NEPC transition and instead performed by SRCAP. The separation of TIP60-C chaperone and acetyltransferase activity upon transition to NEPC raises the intriguing question of whether this potentiates TIP60-C coactivator function via dedicated H2A.Z acetylation, a topic for future investigation.

### NEPC TFs function as a network, culminating in chromatin recruitment of TIP60-C by MYCL

Having documented a critical role of TIP60-C and H2A.Z acetylation in NEPC survival, we returned to the set of NEPC dependency TFs identified earlier (**Figure 2**) to explore whether they function collaboratively and, if so, how that collaboration might connect to TIP60-C. As a first step, we performed a series of CUT&RUN experiments to map the chromatin binding sites of each TF as well as TIP60. The fidelity of the data was confirmed by motif analysis, showing enrichment of the corresponding TF motif for each CUT&RUN assay (**Figure S5A**). Anchoring the analysis on PRAD and NEPC specific and shared cis-regulatory elements (CRE), which were enriched for enhancers and promoters, respectively, as demonstrated by H3K27ac and H3K4me3 occupancy, we observed a striking overall coincidence of TF binding across the entire NEPC enhancer/promoter landscape (**Figure 5A**). Furthermore, TFs expressed in both lineages, such as FOXA1 and NFIB, relocalized from PRAD to NEPC enhancers. ASCL1, FOXA2, SOX1/11, and MYCL, all of which are exclusively expressed in NEPC, also mapped to NEPC enhancers/promoters with a heatmap profile overlapping the sites bound by FOXA1 and NFIB. As expected, all the NEPC TFs broadly overlapped with TIP60, H2AZ and H2AZac, each of which also displayed the same PRAD to NEPC enhancer relocalization seen with FOXA1 and NFIB.

**Figure 5.**
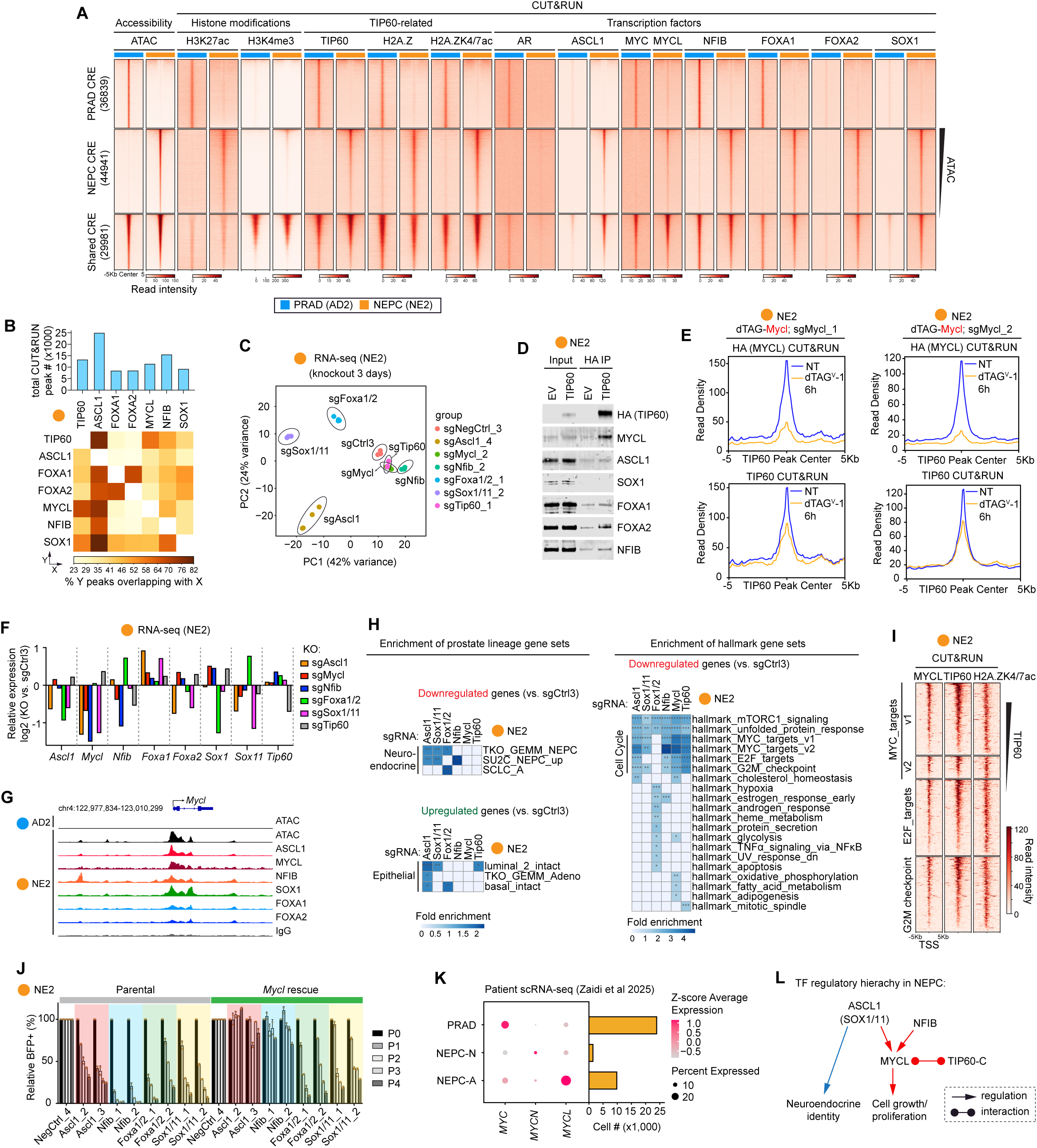
NEPC TFs function as a network, culminating in chromatin recruitment of TIP60-C by MYCL. (**A**) Heatmap showing chromatin accessibility (ATAC-seq) and occupancy (CUT&RUN) of indicated histone modifications and proteins at NEPC-specific, PRAD-specific and shared CREs defined by ATAC-seq in NE2 and AD2 organoids. (**B**) (Top) Total CUT&RUN peak numbers of indicated TFs or TIP60. (Bottom) Heatmap showing percentage of peak overlap through pairwise intersection analysis of indicated TFs and TIP60. (**C**) PCA of RNA-seq of NE2 organoids upon 3-day knockout of indicated gene(s). Double knockout of Foxa1/2 and Sox1/11 were performed. n = 3 independent sgRNA transductions for sgNegCtrl3, sgTip60, sgMycl and sgNfib. n = 4 independent sgRNA transductions for sgAscl1, sgSox1/11, sgFoxa1/2. (**D**) Western blot analyses of immunoprecipitation with HA tag in NE2 organoids expressing HA-FLAG-tagged Tip60 (Tip60 dTr4) or empty vector (EV). (**E**) Average profile showing occupancy (CUT&RUN) of MYCL (via HA tag) and TIP60 at TIP60 peaks in NE2 organoids with expression of dTAG-Mycl and knockout of endogenous *Mycl* using two independent sgRNAs (see **Figure S5F** for heatmaps). Organoids were treated with 500 nM dTAG^V^-1 for 6 h. NT, non-treated. (**F**) Differential expression of TFs and *Tip60* (RNA-seq) upon 3-day knockout of indicated gene(s) compared to control sgRNA in NE2 organoids. (**G**) Genome browser view of chromatin accessibility (ATAC-seq) and occupancy (CUT&RUN) of indicated TFs at the *Mycl* locus in AD2 or NE2 organoids. (**H**) Overrepresentation analyses (FDR < 0.05) of custom neuroendocrine/neuronal signatures (9 sets) and hallmark gene sets (50 sets) in upregulated (FDR < 0.05, log2FC > 0) and downregulated (FDR < 0.05, log2FC < 0) genes upon 3-day knockout of indicated TFs or *Tip60* compared to control sgRNA in NE2 organoids (see also **Figure S5G**). ****, FDR < 0.0001. ***, FDR < 0.001. **, FDR < 0.01. *, FDR < 0.05. (**I**) Heatmap showing occupancy (CUT&RUN) of MYCL, TIP60 and H2A.Zac at the TSS of genes within indicated hallmark gene sets. TSS, transcription start site. (**J**) Cell competition assays using two independent single or dual sgRNA vectors targeting indicated TF or TF paralogs in parental NE2 organoids or those overexpressing *Mycl* (with HA-FLAG-dTAG), showing relative BFP percentage over 4 passages (P0-4). BFP percentages were normalized to day 2 (P0) and the NegCtrl4 sgRNA at the respective day (relative %BFP+). Data represent mean ± SD from n = 2 technical replicates. (**K**) Expression of MYC paralogs in PRAD, NEPC-N and NEPC-A tumor cells in a published scRNA-seq dataset of human prostate tumor biopsies^8^. (**L**) Schematic depicting a TF regulatory hierarchy in NEPC. *Ascl1* acts upstream to 1) promote transcriptional programs underlying neuroendocrine identity and 2) activate *Mycl*, which acts downstream to drive transcriptional programs of cell growth/proliferation.

To more precisely measure TF colocalization, we performed pairwise comparisons of TF binding sites to determine the percentage overlap with each other and with TIP60. ASCL1 is notable for its most broad genomic distribution and high (>70%) overlap with all the other TFs (and with TIP60), consistent with its known role as a master regulator of neural/neuroendocrine lineage specification^65–67^ (**Figure 5B**). In contrast, overlap of FOXA1 and FOXA2 peaks with other factors was more limited (primarily with each other). MYCL was notable in that it had highest cistrome similarity with TIP60 than any of the other TFs (**Figures 5B and S5B)**.

To explore the implications of the striking overlap in TF binding on gene expression, we examined global transcriptomic changes (via RNA-seq) 3 days after knockout of each TF or TIP60, before the antiproliferative consequences of knockout are manifest. In the case of FOXA1/FOXA2 and SOX1/SOX11, both paralogs were co-deleted to ensure complete TF family perturbation. Principal component analysis (PCA) revealed that each TF family has a distinct transcriptomic profile, with the changes incurred following *Ascl1* knockout producing the greatest shift across PC1 and PC2 (**Figure 5C**), again consistent with its master regulator function. Remarkably, *Mycl* and *Tip60* knockout showed near complete overlap in PCA space (**Figure 5C**) and shared the highest number of regulated genes (**Figure S5C**). Because the MYCL paralog MYC (c-MYC) can bind TRRAP^60,61,68^, we postulated that MYCL might also bind TIP60-C for transcriptional coactivation. Indeed, immunoprecipitation experiments revealed that TIP60-C forms a complex with MYCL protein (**Figure 5D**). Because MYCL contains sequence-specific DNA binding activity whereas TIP60 does not, we next asked if MYCL directly recruits TIP60 to its target sites. Using NEPC organoids expressing a *dTAG-Mycl* allele (see methods), we observed a global reduction in TIP60 binding 6 hours after MYCL degradation in two independently derived NEPC tumoroid lines (**Figures 5E and S5D-F**), establishing that MYCL recruits TIP60-C to chromatin through physical interaction.

To determine if any of the TFs within the network regulated the expression of other network TFs, we performed CRISPR knockout of each TF (and *Tip60*) then measured mRNA levels of the remaining TFs 3 days later. *Ascl1* and *Sox1* mRNA levels were substantially reduced following *Foxa1/2* knockout (**Figure 5F**), consistent with recent work showing that FOXA1 acts as an earlier pioneer in the PRAD-to-NEPC transition^69^. Most striking was the reduction in *Mycl* mRNA after knockout of every TF assessed, with the exception of *Foxa1/2* which only affected *Ascl1* and *Sox1* expression (**Figure 5F**). Consistent with these data, CUT&RUN experiments showed that overlapping ASCL1, NFIB and SOX1 binding peaks are seen at the MYCL enhancer/promoter regions (as well as MYCL itself, suggesting autoregulation) (**Figure 5G**). Collectively, the data suggest that ASCL1, NFIB and SOX1 collaborate to directly induce *Mycl* expression, then MYCL subsequently partners with TIP60-C to activate MYCL target genes.

To determine the transcriptional programs regulated by each TF, we performed gene set expression analysis after short term (3 day) knockout of each TF. *Ascl1* deletion led to downregulation of NEPC lineage genes and upregulation of luminal and PRAD lineage programs (**Figure 5H**), consistent with the role of ASCL1 as NE lineage master regulator. Similar changes were seen following combined *Sox1*/*Sox11* knockout, suggesting collaboration of SOX1/11 with ASCL1 (also seen in overlap of TF binding peaks, **Figure 5B**, and coregulated genes, **Figure S5C**). Conversely, deletion of *Mycl* or *Tip60* (as well as *Nfib*) resulted in downregulation of MYC targets, E2F targets and G2M checkpoint gene sets but no changes in lineage identity (**Figures 5H and S5G**). Moreover, CUT&RUN analysis revealed occupancy of MYCL, TIP60 and H2A.Zac at the promoters of genes within such pro-proliferation transcriptional programs (**Figure 5I**), indicating direct regulation. These same gene sets were also downregulated following *Ascl1* or *Sox1*/*Sox11* knockout (but not *Foxa1*/*Foxa2* knockout), consistent with the proposal above that MYCL/TIP60 functions downstream of *Ascl1*. As a functional test of this model, we uncoupled MYCL expression from ASCL1 control in NEPC cells by expressing *Mycl* from a heterologous promoter, then evaluated cell growth following *Ascl1* knockdown. Remarkably, heterologous *Mycl* expression fully rescued the proliferation defect seen with *Ascl1* deletion but not that seen with *Foxa1*/*Foxa2* deletion (**Figure 5J**). Consistent with MYCL acting downstream of ASCL1, *MYCL* expression was markedly upregulated in ASCL1+ human NEPC cells compared to NEUROD1+ NEPC or PRAD, which primarily express MYC (**Figure 5K**)^8^. We conclude that MYCL acts downstream of ASCL1 to drive NEPC proliferation, whereas ASCL1 functions upstream to also drive the NE lineage state (see model, **Figure 5L**).

### MYC paralogs partner with TIP60 for coactivation exclusively in the NEPC lineage of prostate cancer

The fact that MYC pathway plays such a critical downstream role in driving proliferation of NEPC tumoroids raises the broader question of whether the level of MYC activity varies across lineages. Indeed, gene sets for MYC targets, E2F targets and G2/M checkpoint were all substantially upregulated in NEPC tumoroids compared to isogenic wildtype and PRAD tumoroids (**Figure 6A**). MYC targets are also significantly upregulated in human NEPC compared to PRAD (**Figure 6B**). To determine if this increase in MYC activity might lead to greater MYC dependency, we again turned to the dTAG system to investigate the antiproliferative consequences of MYCL degradation in NEPC tumoroids versus MYC degradation in PRAD tumoroids (recall the switch from MYC to MYCL during the PRAD to NEPC transition) (**Figures 6C and S6A)**. Remarkably, NEPC tumoroids were 2-3 orders of magnitude more sensitive to MYCL degradation than PRAD organoids were to MYC degradation, as measured by growth IC50 (**Figure 6D**). Modest (∼50%) degradation of MYCL had profound effects on viability/proliferation in NEPC tumoroids, whereas comparable degradation of MYC in PRAD tumoroids had no discernable consequences. These data reveal a remarkable degree of MYC pathway addiction, similar to results seen earlier with TIP60 degradation and consistent with the model that TIP60 is the critical MYCL coactivator.

**Figure 6.**
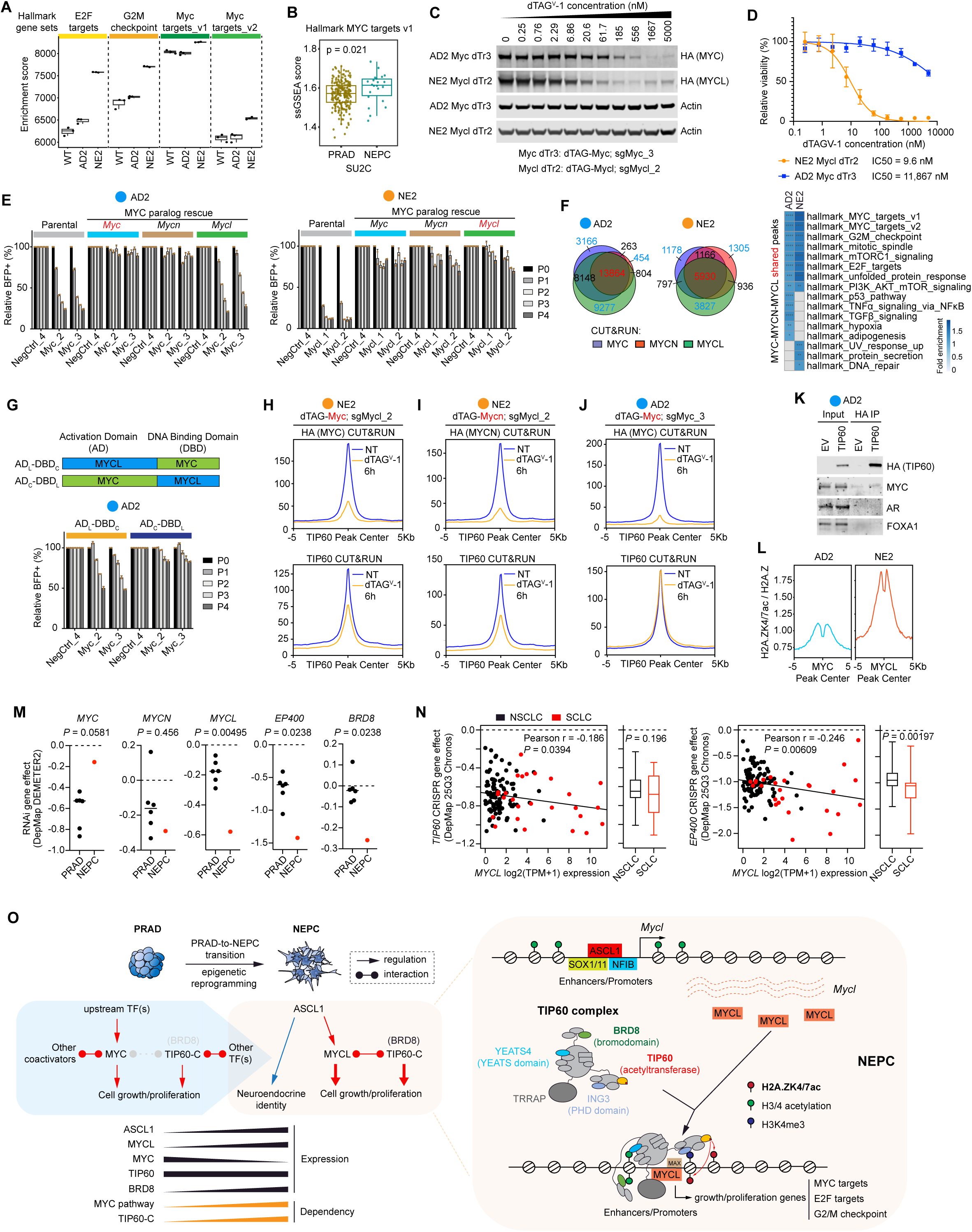
MYC paralogs partner with TIP60 for coactivation exclusively in the NEPC lineage of prostate cancer. (**A**) Boxplot of GSEA enrichment scores (RNA-seq) showing expression of indicated hallmark gene sets in WT, AD2 and NE2 organoids. (**B**) Boxplot of single-sample GSEA (ssGSEA) scores (RNA-seq) showing expression of hallmark MYC target genes in PRAD and NEPC patient tumor samples of the SU2C cohort. (**C**) Western blots of whole cell extract from indicated AD2 dTAG-Myc (dTr3) and NE2 dTAG-Mycl (dTr2) lines treated with a 3-fold gradient of dTAG^V^-1 for 24 hours. Actin was used as loading control. (**D**) dTAG^V^-1 dose response curves and IC50 values for indicated dTAG lines. Tumoroid lines were treated with the same dTAG^V^-1 gradient as in (**C**) for 5 days before CellTiter Glo assays were performed to determine cell viability. Data represent mean ± SD from n = 10 biological replicates. (**E**) Cell competition assays using two independent Mycl sgRNAs in parental AD2 and NE2 organoids or those overexpressing *Myc, Mycn, or Mycl* (with HA-FLAG-dTAG). In (**E**) and (**G**), data represent relative BFP percentage over 4 passages (P0-4). BFP percentages were normalized to day 2 (P0) and the NegCtrl_4 sgRNA at the respective day (relative %BFP+). Data represent mean ± SD from n = 2 technical replicates. (**F**) (Left) Venn diagram showing overlap of CUT&RUN peaks of MYC, MYCN and MYCL (via HA tag) expressed in AD2 and NE2 tumoroids as in (**E**). (Right) Overrepresentation analysis (FDR < 0.05) of hallmark gene sets (50 sets) in genes associated with overlapping CUT&RUN peaks of MYC, MYCN and MYCL in AD2 and NE2 organoids (see **Figure S6C** for analysis with unique peaks). ****, FDR < 0.0001. ***, FDR < 0.001. **, FDR < 0.01. *, FDR < 0.05. (**G**) (Top) Schematic showing the engineered chimeric MYC proteins. (Bottom) Cell competition assays using two independent *Mycl* sgRNAs in AD2 organoids overexpressing chimeric MYC proteins (with HA-FLAG-dTAG). (**H**) Average profile showing occupancy (CUT&RUN) of MYC (via HA tag) and TIP60 in NE2 tumoroids with expression of dTAG-Myc and knockout of endogenous *Mycl* alleles. In (**H**), (**I**), and (**J**), organoids treated with 500 nM dTAG^V^-1 for 6 h. NT, non-treated. (**I**) Average profile showing occupancy (CUT&RUN) of MYCN (via HA tag) and TIP60 in NE2 tumoroids with expression of dTAG-Mycn and knockout of endogenous *Mycl* alleles. (**J**) Average profile showing occupancy (CUT&RUN) of MYC (via HA tag) and TIP60 in AD2 tumoroids with expression of dTAG-Myc and knockout of endogenous *Myc* alleles. (**K**) Western blot analyses of immunoprecipitation with HA tag in AD2 organoids expressing HA-FLAG-tagged Tip60 (Tip60 dTr4) or empty vector (EV). (**L**) Average profile showing CUT&RUN H2A.Zac enrichment (normalized against total H2A.Z levels) at MYC peaks in AD2 and MYCL peaks in NE2 tumoroids. (**M**) Dependency gene effect (RNAi) of indicated MYC paralogs and TIP60-C subunits in PRAD and NEPC lines from the DepMap DEMETER2 dataset. The only NEPC line NCI-H660 is labelled in red. (**N**) Scatter plot depicting the linear association between the *TIP60* (left) and *EP400* (right) dependency gene effect (CRISPR) and *MYCL* gene expression in lung cancer in the DepMap 25Q3 Chronos dataset. NSCLC and SCLC lines are labelled in black and red, respectively. *TIP60* and *EP400* dependency gene effect in NSCLC and SCLC was additionally summarized as boxplots. (**O**) (Left) Model showing molecular events underlying the enhanced MYC pathway (MYCL)/TIP60 dependence following PRAD-to-NEPC transition. NEPC transition is accompanied by activation of master TF ASCL1, MYC to MYCL paralog switch, increased expression and incorporation of BRD8 into TIP60-C, and MYC pathway coactivator switch to TIP60-C, culminating in enhanced MYCL/TIP60 dependence in NEPC for growth/proliferation. TIP60-C, the TIP60 complex. (Right) Model showing MYCL partners with the TIP60 coactivator complex to integrate pro-proliferation oncogenic transcriptional outputs of upstream TFs in NEPC. *Mycl* is a transcriptional target of multiple NEPC TFs. MYCL recruits TIP60-C to chromatin through sequence specific DNA binding, which is stabilized by reader subunits of the complex (e.g. BRD8), to acetylate H2A.Z and activate MYCL target genes.

Because the NEPC transition is accompanied by a switch from MYC to MYCL expression, it is possible that the profound increase in MYCL dependency is a consequence of differences in MYC versus MYCL paralogs. To address this question, we performed a series of rescue experiments to ask whether MYC (or MYCN) can rescue the antiproliferative consequences of *Mycl* deletion in NEPC tumoroids. In parallel, we performed the reciprocal experiment to ask if MYCL (or MYCN) can rescue the antiproliferative consequences of *Myc* deletion in PRAD tumoroids. MYC and MYCN were both able to fully rescue NEPC growth at levels comparable to that seen with MYCL, indicating that all three MYC paralogs function similarly to control proliferation (**Figures 6E and S6B**). This conclusion is further supported by CUT&RUN experiments showing high overlap of MYC, MYCN and MYCL chromatin binding peaks in such NEPC tumoroids where endogenous *Mycl* was substituted (**Figure 6F**). In addition, ontology analysis of the genes that map to these shared peaks identified MYC targets, E2F targets and G2/M checkpoint as top hits (**Figure 6F**), further supporting that all three MYC paralogs can directly regulate these key pro-proliferation transcriptional programs.

Surprisingly, the comparable experiment in PRAD tumoroids revealed that MYC and MYCN both fully rescued the proliferation defect caused by *Myc* deletion whereas MYCL did not (**Figure 6E**) despite robust MYCL protein expression (**Figure S6B**). This failure to rescue in PRAD is unlikely explained by differences in DNA binding between MYCL and other MYC paralogs, because, as with NEPC, we observed significant overlap in the chromatin binding peaks of all three paralogs, which also map to MYC targets, E2F targets and G2/M checkpoint hallmark gene sets (**Figure 6F**). This result suggests that the failure of MYCL to rescue PRAD growth may be a consequence of failed coactivation of target genes, perhaps due to differences in N-terminal activation domain, which contains MYC-homology boxes that define the MYC TF family and shows varying degrees of divergence across paralogs^70^ (**Figure S6D**). To test this hypothesis, we performed a domain swapping experiment in which we replaced the activation domain of MYC with that of MYCL and the activation domain of MYCL with that of MYC (**Figures 6G and S6E**). Comparative analysis of these chimeric alleles revealed that PRAD proliferation was restored when the MYC activation domain was coupled with the MYCL DNA binding domain but not when MYCL activation domain was coupled with the MYC DNA binding domain (**Figure 6G**).

Prior work has shown that MYC-homology boxes play a critical role in protein-protein interactions, particularly with coactivators^70–72^. Having shown that the activation domain of MYCL is the reason for failure to rescue PRAD proliferation following MYC knockdown, we postulated this may be due to differences in coactivator usage by different MYC paralog. Earlier we demonstrated that partnership between MYCL and TIP60 is critical for coactivation in NEPC (**Figures 5C-E**). Here we re-examined this question, but now with MYC and MYCN in both PRAD and NEPC lineage background to search for potential paralog- and/or lineage-specific differences. As expected, we found that both MYC and MYCN are critical for chromatin binding of TIP60 in NEPC tumoroids following MYCL depletion. Specifically, chemical degradation of either MYC or MYCN led to a rapid (6 hours) decrease in TIP60 binding (by CUT&RUN) (**Figures 6H**, **6I, S6F, and S6G**), analogous to that seen earlier with MYCL degradation (**Figure 5E**). Conversely, there was no change in TIP60 chromatin binding following MYC degradation in PRAD tumoroids (**Figure 6J**), suggesting that MYC does not partner with TIP60 for coactivation in the PRAD lineage. This conclusion is further supported by the absence of a TIP60-MYC complex in immunoprecipitation experiments in PRAD tumoroids (**Figure 6K**) and a robust increase in H2A.Z acetylation at MYCL peaks in NEPC tumoroids compared to MYC peaks in PRAD tumoroids (**Figure 6L**). In summary, the paralog switch from MYC to MYCL during the PRAD to NEPC transition is accompanied by a switch to TIP60 as the primary MYCL coactivation partner, with increased reliance on H2A.Z acetylation (a key TIP60 substrate) for downstream gene expression. Importantly, the robust partnership between MYCL and TIP60 is not specific to MYCL (based on rescue experiments with MYC and MYCN in NEPC) but is instead defined by the NEPC lineage and results in a greatly enhanced state of MYCL/TIP60 dependency.

As discussed earlier, the dependency on MYCL/TIP60 seen in our murine NEPC models is also evident in human NEPC models such as MSKPCa14 and LuCaP49 (**Figures 2H** and **3H**). To address whether MYCL/TIP60 dependency extends more broadly, we examined the human DepMap database, focusing primarily on human NE cancers. Prostate cancer is poorly represented in DepMap due to the limited number of available cell lines, but it is noteworthy that NCI-H660, the only human NEPC line with perturbation (RNAi) data present in DepMap, shows exquisite MYCL dependence as well as dependence on TIP60-C subunits Ep400 and BRD8 (**Figure 6M**). Conversely, human small cell lung cancer (SCLC) cell lines are abundantly represented and share many transcriptional features and TF dependencies seen in NEPC ^6–8^. TIP60-C dependency (measured by either *TIP60* or *EP400* deletion) is significantly correlated with MYCL expression across all lung cancer cell lines (p=0.0394; p=0.00609, respectively), with the signal largely explained by SCLC lines which express high levels of MYCL, a MYC paralog identified in SCLC^73^ (**Figure 6N**). Dependency scores for *TIP60* and *EP400* sgRNAs are also greater in SCLC lines compared to non-small cell lung cancer (NSCLC), providing further evidence that MYCL/TIP60 dependency may extend broadly to human NEPC and SCLC.

## Discussion

Over the past decade, PRAD to NEPC lineage transition has emerged as an increasingly important mechanism of resistance to ARSI therapy. While TFs such as ASCL1, FOXA2, SOX2 and others have each been implicated in this transition^3–5,18–27,74^, there is limited insight into whether these TFs function independently or collectively to drive NEPC lineage transition. By leveraging a set of isogenic murine tumoroid models that accurately mirror the NEPC transition in patients and can be serially expanded for comprehensive biochemical analysis and functional screens, the following observations emerge. First, through epigenomic analyses we confirm that extensive changes in enhancers largely define the remodeled chromatin landscape seen in NEPC, as expected. Second, through a comprehensive review of the specific TFs potentially driving and/or impacted by the changes in these enhancers, we identify members of 5 TF families (FOXA1/2, ASCL1, SOX1/11, NFIB, MYC) that function as a network to specify and maintain the NEPC lineage. Third, by screening for potential coactivators utilized by these TFs, we identify a critical dependency on the TIP60/KAT5 histone acetyltransferase, which partners with MYCL to drive NEPC proliferation through acetylation of the histone variant H2A.Z. Fourth, through short term TF perturbation experiments, we show that ASCL1 functions upstream to specify neuroendocrine lineage programs and activate MYCL expression. MYCL then acts downstream as the key mediator of cell growth (**Figure 6O**). Finally, through further exploration of MYC family TFs across lineages, we find that the paralog shift from MYC expression in PRAD to MYCL expression in NEPC is accompanied by an unexpected switch to TIP60-C as the principal MYC paralog coactivator in the NEPC lineage state.

While the genetically defined mouse systems used here provide a powerful platform for experimental perturbation, it is also critical that emerging mechanistic insights have relevance to the NEPC clinical setting. We have leveraged a growing panel of PDOs derived from CRPC patients to validate many of the dependencies discovered in the mouse screens, as well as DepMap to extend the findings to human SCLC. One caveat is the mouse system used here accurately models ASCL1+ human NEPC but the relevance to NEUROD1+ human NEPC remains unclear. To that point, one of the NEPC PDOs screened here (MSKPCa10) is NEUROD1 and MYC dependent, yet exquisitely TIP60 dependent (**Figures 2H and 3J**). This magnitude of MYC/TIP60 dependence (versus MYCL/TIP60 dependence in ASCL1+ mouse tumoroids and the human PDO LuCaP49) may reflect broad TIP60 dependence in the NEPC lineage state regardless of which NE-specifying TF (ASCL1 or NEUROD1) or MYC paralog defines that state, as suggested by the MYC paralog rescue experiments in mouse NEPC tumoroids (**Figure 6E**). Although MYCL is the primary focus of the work reported here, it is important to note that MYCN is also a critical MYC paralog in human NEPC^29,75–78^, as evident in some of the isogenic mouse NEPC tumoroids and in human NEPC PDOs (**Figures 2D and 2H**). Furthermore, expression data from human CRPC cohorts confirms a paralog switch from MYC in PRAD to either MYCL or MYCN in NEPC (**Figures 2F**, **5K, and S2F**).

At least three observations from this work raise new questions that warrant further investigation. First, the switch in MYC coactivator usage to TIP60 during the NEPC transition begs the question of which coactivator does MYC utilize in the PRAD lineage state. The MYC/MYCL domain swapping experiments make it clear that the N-terminus of MYCL is unable to access this coactivator in PRAD, providing a potential experimental approach to selectively probe for candidates through pull down studies. Structurally, the absence of MYC-homology box IIIa (MBIIIa) in MYCL (but present in MYC and MYCN) may provide a clue (**Figure S6D**). Interestingly, a known MYC coactivation partner GCN5/PCAF (acetyltransferases of the SAGA coactivator complex)^70,71,79^ has been reported to interact with MYC in part through MBIIIa^61^. GCN5/PCAF deletion scores in our coactivator dependency screen with a trend toward greater dependency in PRAD versus NEPC (**Figure 3A**). Regardless of the answer, it will also be of interest to understand the molecular events that trigger the switch to TIP60 as this might be an opportunity for therapeutic intervention.

A second series of questions center around the mechanism of increased MYCL dependency in the NE lineage state. More specifically, why do NE cells require a much higher threshold of MYC protein for proliferation? Prior work implicating MYC as a transcriptional amplifier of gene expression^80,81^ and reported transcriptional addictions in SCLC and neuroblastoma^82–84^ are consistent with a threshold model; however, there are conflicting reports on the generalizability of the amplifier hypothesis^85^. Another consideration is the established role of MYC in metabolic control^72,86^, raising the possibility that higher levels of MYCL are needed to meet increased metabolic demand, particularly for glucose metabolism (**Figure S6H**), in the NE lineage state. Finally, the increased demand for MYCL and MYC pathway-addiction could be a consequence of switching to TIP60 as the primary coactivator in the NE state, with subsequent reliance on the TIP60 catalyzed H2A.Z acetylation for proliferation/cell cycle gene expression^87,88^. Interestingly, TIP60 and H2A.Z acetylation have been previously implicated in neuronal fate specification^89^. It is also worth noting that, although MYC and TIP60 both score as dependencies in the PRAD state, they function independently. The subsequent biochemical coupling of MYCL and TIP60 as a single TF/coactivator complex during the NE lineage transition could perhaps be linked to the enhanced dependency.

Third, although context-specific subunit exchange has been documented for chromatin complexes such as BAF^90^, TIP60-C assembly has largely been considered static based on biochemical studies in model cell lines. Our observation that BRD8 is selectively (and functionally) incorporated into TIP60-C in NE lineage cells raises the possibility that TIP60-C subunit composition may be dynamically regulated across other cell contexts.

In closing, the high threshold of MYCL dependence seen in NEPC coupled with the switch to TIP60 for coactivation may provide an opportunity to develop novel therapies for NEPC and potentially SCLC. Knockout studies of the various TIP60-C subunits identified two druggable vulnerabilities: (i) the catalytic domain of the TIP60 acetyltransferase and (ii) the reader domain containing proteins BRD8 and YEATS4. Of these, bromodomain protein BRD8 is compelling because it is selectively incorporated into TIP60-C in NEPC, coupled with selective loss of H2A.Z acetylation in NE lineage cells following BRD8 knockout (**Figures 4K and 4L**). Of course, a major caveat with either of these strategies is the potential toxicity associated with pharmacological targeting of a common essential gene implicated in multiple nuclear processes^49^. Nonetheless, the dTAG studies showing a >150-fold difference in sensitivity to TIP60 degradation provide optimism for a potential therapeutic window.

## Acknowledgements.

The authors thank members of the Sawyers laboratory for advice and discussions; Emily Bernstein (Icahn School of Medicine at Mount Sinai), Yu Chen (Memorial Sloan Kettering Cancer Center), Ari Firestone (Calico Life Science LLC) and Cynthia Jung for critical reading of the manuscript; Yu Chen for sharing PDOs; Emily Bernstein for sharing iDFs; Ninghui Mao and Jye Lawson for administrative support; and the High Performance Computing Center, the Molecular Cytology Core, the Antitumor Assessment Core and mouse facilities at MSKCC. Z.S. was supported by an Edith C. Blum Foundation Postdoctoral Training Award. J.L.Z. was supported by the American Society of Clinical Oncology Young Investigator Award, the Prostate Cancer Foundation Young Investigator Award, and a Career Development Award in Clinical Oncology from the National Cancer Institute. T.H. was supported by the Prostate Cancer Research Program Early Investigator Research Award (W81XWH-21-PCRP-EIRA) from the Department of Defense and the Translational Research in Oncology Training (TROT) Program at MSKCC. C.L.S was supported by the Howard Hughes Medical Institute, Calico Life Sciences LLC, and NIH grants CA193837, CA092629, CA265768, CA008748.

## Author Contributions.

Z.S. and C.L.S. conceived of this study. Z.S. and C.L.S. designed experiments and interpreted results. Z.S., J.L.Z., Z.F.K, T.H., S.Y., and M.L. performed experiments. Z.S., W.M.I., S.N., P.C. and R.K. performed computational analysis. N.S. oversaw analyses of patient data. A.G.M. oversaw ChromoHMM analysis and RNA-seq pathway analysis in mouse tumoroids. Z.S. and C.L.S. wrote the manuscript with input from all authors.

## Methods

### Ethical statement

All mouse experiments were conducted under protocol 06-07-012 approved by the Memorial Sloan Kettering Cancer Center (MSKCC) Institutional Animal Care and Use committee (IACUC).

### Organoid derivation and culture

Murine prostate organoids and tumor organoids (tumoroids) were established and maintained in conditions as described previously^92,93^. Human patient derived organoids (PDOs) were cultured as described previously^93,94^. Briefly, for derivation of murine organoids/tumoroids, whole mouse prostates including all lobes or subcutaneous tumors were dissected and minced. Prostates and tumor fragments were digested with collagenase type II (Gibco) for 2 hours at 37 °C, followed by digestion with TrypLE (Gibco) at 37 °C for 1 hour. All digestions were supplemented with Y-27632 (10 μM). The resulting cell suspension was filtered through 40-μm strainers to remove debris. Prostate epithelial cells or tumor cells were embedded in growth factor-reduced Matrigel (Corning) and overlaid with mouse prostate organoid medium. For passaging, organoids were dissociated via TrypLE digestion at 37 °C and pipette trituration through 200-μl tips, except mouse NEPC tumoroids, which were dissociated directly via pipette trituration. Dissociated organoids were embedded in 50-μl drops of growth factor-reduced Matrigel at 7,000 – 30,000 cells per drop and overlaid with mouse or human prostate organoid medium. To harvest NEPC tumoroids for lentiviral transduction, cell sorting, and molecular/biochemical analysis, tumoroids were incubated with Cell Recovery Solution (Corning) on ice for 30 min to dissolve Matrigel.

### Establishment of isogenic PRAD and NEPC organoid series

*Pten*, *Rb1*, *Trp53* TKO PRAD and NEPC tumors were derived using an organoid transplantation-based tumor formation and lineage transformation system^3,28^. Prostates of *Rosa26*^mT/mG^*Rb1^fl/fl^Trp53^fl/fl^Pten^fl/fl^*mice^30^ (without *PBCre*) were harvested from adult mice and dissociated. Cells were stained on ice for 1 hour with lineage marker cocktail (Lin), EpCAM, CD49F, CD24A, CD52, Sca1, and Prom1 antibodies. Luminal 2 (L2) prostate epithelial cells (Lin^-^/EpCAM^+^/CD24A^+^/CD49F^-^/CD52^-^/Sca1^+^)^91^, which we observed have highest efficiency for NEPC transition compared to luminal 1 (L1) or basal cells, were isolated via fluorescence-activated cell soring (FACS) and utilized to establish normal prostate organoid (WT) (7-10 days). To induce Cre recombinase expression, adenoviral transduction was performed by mixing normal prostate organoids with Ad-CMV-iCre (Vector Biolabs) and spinoculation to simultaneously induce deletion of the floxed tumor suppressors and activate GFP expression; GFP^+^ cells (*Rb1^-/-^Trp53^-/-^Pten^-/-^*) were purified via FACS to establish TKO organoids (TKO-AD1) (7-10 days). TKO organoids were further transplanted subcutaneously into NSG mice (JAX) to allow tumor development. Briefly, 500,000 organoid cells were resuspended in 100 μl of a 1:1 mixed Matrigel and organoid culture medium and injected subcutaneously into the right flank of immunodeficient NSG mice at 2 months of age. Tumors were harvested after 4-6 months for histological analysis and tumoroid derivation. Tumoroids from TKO tumors were further screened for PRAD and NEPC histological phenotypes and tumor marker expression by immunofluorescence and RT-qPCR (RNA). We further enriched for homogeneous PRAD or NEPC phenotype. Tumoroid lines with high PRAD or NEPC histology/marker expression were further subcloned by manually picking individual organoids under a dissection microscope, followed by serial passage and expansion in organoid culture. TKO-AD2 (PRAD), TKO-NE1 (NEPC) and TKO-NE2 (NEPC) were established as stable, pure PRAD or NEPC tumoroid lines.

### Histology and immunostaining

Organoids embedded in Matrigel were fixed using 4% paraformaldehyde, dehydrated with 70% ethanol, paraffin-embedded and sectioned. H&E staining was performed following standard protocols by the MSKCC Molecular Cytology Core.

Immunofluorescence were performed on a Leica Bond RX automatic stainer using antibodies listed in **Supplementary Table 1**. All formalin-fixed paraffin-embedded stained tissue was scanned using a MIRAX scanner.

### Intracellular flow cytometry

Intracellular staining of DLL3 was performed using a PE-conjugated DLL3 antibody (BioLegend, 154004) and the BD Cytofix/Cytoperm Fixation/Permeabilization Kit (BD Biosciences) as previously described^14^.

### Plasmids and sequences

For construction of expression vectors, all sequences, including cDNA, epitope tag (2xHA and 3xFLAG), degron tag (dTAG), and fluorescent reporter (mCherry or GFP), were ordered as gene fragments from Twist Bioscience. DNA fragments were assembled into a pRRL-SFFV lentiviral vector using In-Fusion cloning kit (Takara). Epitope and degron tags were in-framed fused to the N-terminus of target proteins and fluorescent reporters were added to the C-terminal side following a P2A or IRES sequence. For cloning of CRISPR sgRNA (knockout and CRISPRi), oligos ordered from IDT were annealed in T4 ligation buffer (NEB) and phosphorylated using T4 PNK (NEB). The annealed oligos and BsbBI (NEB)-linearized pLV-hU6-sgRNA-SFFV-Puro-P2A-BFP plasmid were ligated using T4 DNA ligase (NEB). The pLV-hU6-sgRNA-SFFV-Puro-P2A-BFP vector backbone was modified from a published hU6-sgRNA-SFFV-Puro-P2A-GFP vector^95^ by replacing GFP with TagBFP. For construction of dual sgRNA vectors, sequences containing the first sgRNA, mU6 promoter and the second sgRNA were ordered as gene fragments from Twist Bioscience and assembled into BsbBI (NEB)-linearized pLV-hU6-sgRNA-SFFV-Puro-P2A-BFP plasmid, resulting in pLV-hU6-sgRNA-mU6-sgRNA-SFFV-Puro-P2A-BFP. sgRNAs of high specificity and efficiency were selected from the ‘CRISPR 10K’ track of the UCSC genome browser. Sequences of sgRNAs used in this study are listed in **Supplementary Table 2**.

### Organoid engineering

For lentiviral transduction, 50,000 – 200,000 dissociated organoid cells were spin-infected with lentivirus mixed with 8 μg/ml Polybrene (Millipore Sigma) at 600 xg and 32 °C for 1 hour followed by an additional 30 min incubation in the tissue culture incubator. Transduced cells were then seeded into organoid culture. To establish organoid lines with stable transgenic expression, dissociated organoids cells were subject to lentiviral transduction, followed by cell sorting (FACS) using a SONY MA900 instrument with 100um sorting chips or antibiotic selection to enrich for cells containing the lentiviral construct which encodes a fluorescent marker or antibiotic resistance gene. Organoids transduced with lentiviral Cas9 (lenti-Cas9-blast) or dCas9-KRAB (lenti-dCas9-KRAB-blast) were selected using blasticidin (10 μg/ml). Genetic knockout was performed using CRISPR-Cas9 technology. Organoids stably expressing Cas9 was infected with lentiviral sgRNA (pLV-hU6-sgRNA-SFFV-Puro-P2A-BFP) or dual sgRNA (pLV-hU6-sgRNA-mU6-sgRNA-SFFV-Puro-P2A-BFP). Knockout cells were further purified when needed using cell sorting for BFP^+^ cells or puromycin (2 μg/ml) selection (NE1 and NE2 murine tumoroids were selected using 0.2 μg/ml puromycin due to high sensitivity). For CRISPR interference (CRISPRi), organoids stably expressing dCas9-KRAB was infected with lentiviral sgRNA. Knockdown cells were further purified when needed using puromycin selection.

### Competition-based proliferation assay and screen

Cas9-expressing organoid cells were transduced with lentiviral sgRNA (pLV-hU6-sgRNA-SFFV-Puro-P2A-BFP). Baseline BFP percentages as a readout of infection rate were determined by flow cytometric analysis using a BD LSRFortessa instrument at day 2 post transduction (P0) before any growth defect manifest. Typically, 40-70% BFP^+^ cells were observed at baseline. BFP percentages were measured every 4 (WT, AD2 and NE2 murine organoids and PCa2, PCa3, PCa11, PCa12 PDOs) or 7 (NE1 murine organoids and PCa10, PCa14 and LuCaP49 PDOs) days during passage and tracked over 4 passages (P1 to P4). BFP percentages of each sgRNA over the time course were normalized to the initial percentage (P0) and that of Negative Control sgRNA (sgNegCtrl) at the corresponding day. 2 to 4 independent sgRNAs targeting DNA binding domains of TFs, catalytic domains of coactivator enzymes, and reader domains of chromatin reader proteins, when applied, were used for each gene. Screens were performed in a well-by-well format where organoids were transduced with individual sgRNA(s). For TF screens, the average percentage of BFP+ cells of all sgRNAs used was used to determine sgRNA-containing cell population.

### Establishment of dTAG organoids

Organoids expressing Cas9 were transduced with lentiviral constructs encoding N-terminally 2xHA-3xFLAG-dTAG tagged *Tip60*, *Myc*, *Mycn*, or *Mycl* linked with mCherry (GFP containing version was used only for expression in WT organoids). Transgenes were engineered to contain same sense mutation(s) at PAM or seed sequences to confer resistance to sgRNA targeting. Organoids were dissociated 3 days post transduction and transduced cells were purified using FACS. Organoids were expanded and further transduced with lentiviral sgRNA constructs (pLV-hU6-sgRNA-SFFV-Puro-P2A-BFP) followed by puromycin selection. BFP percentage was measured by flow cytometric analysis to ensure purity of population. Expression of dTAG transgenes and knockout of endogenous genes were further confirmed by western blotting. Stable organoid lines were maintained in culture supplemented with blasticidin and puromycin.

### Growth and viability assay

Growth assays were performed using CellTiter Glo (Promega) following manufacture’s protocol. Briefly, dissociated organoids were embedded in a 30-μl drop in each well of round bottom 96-well dish and seeded in 10 replicates. Organoids were treated with indicated concentrations of dTAG^V^-1 (Tocris), AU-15330 (MedChemExpress), BAY-876 (MedChemExpress), or V-9032 (MedChemExpress) for 6 (mouse organoids) or 10 – 13 (PDOs) days before viability was measured using the CellTiter Glo assay with a GloMax Navigator instrument (Promega).

### Whole cell extract preparation and western blotting

Whole cell extract was prepared as previously described^96^. Briefly, dissociated organoid cells were lysed in TOPEX buffer (50 mM Tris-HCl pH 7.5, 300 mM NaCl, 1 mM MgCl_2_, 0.5% Triton X-100, 1% SDS, 1 mM DTT, 1x protease inhibitor cocktail (MilliporeSigma) and 125 U/ml Benzonase (MilliporeSigma)) at room temperature for 10 min and protein concentration was determined by BCA assay (Thermo Fisher). Samples were boiled in Laemmli buffer (Bio-rad) at 70 °C for 10 min for immunoblotting following standard protocols using antibodies listed in **Supplementary Table 1**. Proteins were visualized via fluorescence detection using the Odyssey CLx imaging system (LI-COR Biosciences) or using SuperSignal West Pico PLUS Chemiluminescent Substrate (Pierce) and the Amersham ImageQuant 800 imaging system (GE healthcare).

### Immunoprecipitation

Immunoprecipitation was performed as previously described^39^ with modifications. For crude nuclei isolation, 10 – 20 million dissociated organoid cells were mixed with 1 ml of Hypotonic Buffer (10 mM HEPES, pH 7.9, 10 mM KCl, 1.5 mM MgCl_2_, 1 mM DTT, and 1x protease inhibitor cocktail) followed by pipette trituration. Nuclei were extracted for 40 min for NE2 organoids and 2 hours for AD2 and WT organoids at 4 °C with 500 μl High Salt Buffer (20 mM HEPES, pH 7.9, 420 mM NaCl, 1.5 mM MgCl_2_, 0.2 mM EDTA, 25% glycerol, 1 mM DTT, and 1x protease inhibitor cocktail) supplemented with 0.1% NP-40 and 125 U/ml Benzonase. Nuclei were pelleted and supernatant taken for immunoprecipitation. The nuclear extract was diluted with 1.8 volumes (900 μl) of Low Salt Buffer (20 mM HEPES, pH 7.9, 1.5 mM MgCl_2_, 0.2 mM EDTA, 20% glycerol, 1 mM DTT, and 1x protease inhibitor cocktail) supplemented with 0.1% NP-40 to obtain a final NaCl concentration of 150 mM, and cleared at 40,000g for 30 min at 4 °C. Immunoprecipitation was performed with 25 μl of HA-trap Magnetic Agarose beads (ChromoTek) overnight at 4 °C. The beads were then washed twice with 1 ml Buffer G 150 (50 mM Tris, pH 7.5, 150 mM NaCl, 0.5% NP-40), twice with 1 ml Buffer G 250 (50 mM Tris, pH 7.5, 250 mM NaCl, 0.5% NP-40) and boiled in Laemmli buffer at 70 °C for 10 min for immunoblotting using antibodies listed in **Supplementary Table 1**.

### RNA extraction, cDNA synthesis, and RT-qPCR

Total RNA was extracted using RNeasy Plus Mini Kit (Qiagen) according to manufacturer’s protocol. 1 mg of total RNA was subjected to reverse transcription using First-Strand cDNA Synthesis System (ORIGENE). qPCR was performed using PowerUp SYBR Green Master Mix (Applied BioSystems) and the QuantStudio 6 Flex Real-Time PCR System (Applied BioSystems) with primers listed in **Supplementary Table 3**.

### RNA-seq

RNA-seq was performed in triplicates of independent cultures. For short-term knockout experiments, BFP+ sgRNA-containing cells were sorted out using a SONY MA900 instrument 3 days after lentiviral delivery of sgRNA vectors. Total RNA was extracted using RNeasy Plus Mini Kit (Qiagen) according to manufacturer’s protocol. 100 – 1000 ng of total RNA was used for poly(A) RNA selection using the NEBNext Poly(A) mRNA Magnetic Isolation Module (NEB). Strand-specific RNA libraries were prepared using the NEBNext Ultra II Directional RNA Library Prep Kit for Illumina (NEB) according to manufacturer’s protocol. Quality of libraries was analyzed using Qubit fluorometer and Agilent TapeStation. Barcoded libraries were multiplexed and subjected to 75-bp single-end sequencing with an Illumina NextSeq 550 instrument.

### ATAC-seq

ATAC-seq was performed as previously described^96^ in duplicates of independent cultures. 50,000 cells were washed once with 50 μl cold PBS in tubes pre-coated with 1% BSA. Cells were lysed in 50 μl cold Lysis Buffer (10 mM Tris-HCl pH 7.4, 10 mM NaCl, 3 mM MgCl_2_, 0.1% Tween-20, 0.1% NP-40, and 0.01% Digitonin) for 5 min on ice. Lysis was stopped by adding 1 ml of cold Wash Buffer (10 mM Tris-HCl pH 7.4, 10 mM NaCl, 3 mM MgCl_2_, and 0.1% Tween-20) and nuclei were spun down by centrifuging at 600 xg for 10 min at 4 °C. Nuclei pellet was resuspended in 50 μl cold Transposition Mix (25 μl 2x TD buffer (Illumina), 2.5 μl Tn5 transposase (Illumina), 16.5 μl PBS, 0.5 μl 1% Digitonin, 0.5 μl 10% Tween-20, and 5 μl H2O) and the reaction was incubated at 37 °C for 30 min in a thermomixer with 1000 rpm mixing. Transposed genomic DNA was purified using a Zymo DNA Clean and Concentrator-5 Kit and subjected to PCR amplification using NEBNext High-Fidelity 2x PCR Master Mix with custom Nextera PCR primers. Optimal PCR cycles were determined by qPCR using partially amplified libraries. Libraries were cleaned up and size-selected using double-sided bead purification with Agencourt AMPure XP magnetic beads (Beckman Coulter) to remove primer dimers and fragments longer than 1 kb. Library quality control was performed with Qubit fluorometer and TapeStation. Barcoded libraries were multiplexed and subjected to 35-bp paired-end sequencing with an Illumina NextSeq 550 instrument.

### CUT&RUN

CUT&RUN was performed as previously described^96^. Briefly, 0.5 x 10^6^ cells were washed three times with Wash Buffer (20 mM HEPES-NaOH pH 7.5, 150 mM NaCl, 0.5 mM Spermidine, 1x protease inhibitor cocktail) at room temperature. Cells were resuspended in Wash Buffer, mixed with activated BioMag Plus Concanavalin A (Con A) beads and incubated for 10 min at room temperature. Cell-Con A bead conjugates were mixed with 50 μl cold Antibody Buffer (20 mM HEPES-NaOH pH 7.5, 150 mM NaCl, 0.5 mM Spermidine, 0.01% Digitonin, 2 mM EDTA, 1x protease inhibitor cocktail) with antibodies listed in **Supplementary Table 1** and incubated overnight at 4 °C on a nutator. Beads were washed three times with cold Digitonin Buffer (20 mM HEPES-NaOH pH 7.5, 150 mM NaCl, 0.5 mM Spermidine, 0.01% Digitonin, 1x protease inhibitor cocktail) at 4 °C. Beads were then resuspended with 50 μl cold Digitonin Buffer and pAG-MNase^96^ was added at a 1:1000 ratio, followed by incubation at 4 °C for 1 h on a nutator. Beads were washed three times with cold Digitonin Buffer at 4 °C. Beads were resuspended with 50 μl cold Digitonin Buffer and pAG-MNase was activated by adding CaCl_2_ to 2 mM. Targeted chromatin cleavage was carried out by incubation at 4 °C for 30 min on a nutator. The reaction was stopped by adding 50 μl cold 2x Stop Buffer (340 mM NaCl, 20 mM EDTA, 4 mM EGTA, 0.01% Digitonin, 50 mg/ml RNase A, 1 pg/ul Drosophila spike-in DNA). Cleaved chromatin was released by incubating at 37 °C for 30 min in a thermomixer with 750 rpm mixing. Beads were magnetically collected and CUT&RUN DNA (> 50 bp) was extracted from the supernatant using the Monarch DNA Cleanup Kit (NEB). CUT&RUN library was prepared with no more than 10 ng CUT&RUN DNA using NEBNext Ultra II DNA Library Prep Kit (NEB) according to manufacturer’s protocol with modifications: end prep was performed for 30 min at 20 °C followed by 1 h at 50 °C, adaptor-ligated DNA was cleaned up without size selection, and the anneal/extension step of PCR amplification was performed for 10 s at 65 °C. Library quality control was performed with Qubit fluorometer and TapeStation. Barcoded libraries were multiplexed and subjected to 35-bp paired-end sequencing with an Illumina NextSeq 550 instrument.

### RNA-seq analysis

Raw bulk RNA-seq data were processed using bcl2fastq (v2.20.0.422, Illumina) and quality checked using FastQC (v0.11.8) (https://www.bioinformatics.babraham. ac.uk/projects/fastqc/). Single-end sequencing reads were aligned to the mouse genome (GRCm38.p6/mm10) using STAR (v2.6.0a)^97^ with default parameters. The number of mapped reads was counted using STAR with ‘‘–quantMode GeneCounts’’. Heatmaps of z-score transformed expression values and principal component analysis (PCA) plots were generated using DESeq2 (v1.30.0)^98^. Differential gene expression between each group was performed using DESeq2 with default settings, where the counts are fitted to a negative binomial generalized linear model followed by Wald test for significance and log fold change shrinkage using apeglm shrinkage estimator. Genes with false discovery rate (FDR) < 0.05 were considered differentially expressed with log2 fold change > 0 as upregulated and log2 fold change < 0 as downregulated. Upset plots were made using UpSetR^99^ and ComplexHeatmap ^100^ R packages. Gene set over-representation analysis (ORA) was performed using the enricher function (hypergeometric test with Benjamini-Hochberg correction for multiple testing) and gene set enrichment analysis (GSEA) was performed using the GSEA function, both from the clusterProfiler R package^101^. Hallmark gene sets (50 sets) were obtained from the msigdbr database. Prostate lineage gene sets from GEMM^7,91^ and patients (SU2C^31^ and WCM^29^) that were published or curated in this study were listed in **Supplementary Table 4**. For both ORA and GSEA, FDR < 0.05 is considered statistically significant. Bulk RNA-seq data of MSKCC series of patient derived organoids (PDOs) were obtained from GEO (GSE199190). LuCaP49 patient (PDX) derived organoids^7^ RNA-seq data was generated in this study. RNA-seq data of all PDOs were re-analyzed as described in this section.

For analysis on bulk RNA-seq of human prostate cancer, data from two clinical cohorts were obtained from the cBioPortal for Cancer Genomics^102^. The cohorts included: (1) the SU2C/PCF Dream Team cohort (“SU2C”, 210 PRAD and 22 NEPC samples)^31^, and (2) the Beltran et al. cohort (“WCM”, 34 PRAD and 15 NEPC samples)^29^. Hallmark MYC signature scores from the Molecular Signatures Database (MSigDB) were calculated using the Gene Set Variation Analysis (GSVA) R package^103^ with the single-sample gene set enrichment analysis (ssGSEA) method. Statistical differences in signature scores and mRNA expression for individual genes of interest between PRAD and NEPC samples were evaluated using a two-tailed Wilcoxon rank-sum test.

For analysis on scRNA-seq of human prostate cancer, data from the Zaidi et al. cohort (35,696 cells from 23 tumor samples)^8^ were obtained from GEO (GSE264573). The count matrix was normalized by (1) dividing feature counts by total feature counts per cell, (2) scaling by a factor of 10,000, and (3) applying natural-log transformation with a pseudo-count of 1 (’NormalizeData’). Cell-cycle signatures were subsequently scored (’CellCycleScoring’) and the resulting scores were regressed out prior to feature scaling and centering (’ScaleData’). Cell annotations from the original publication were retained. Specifically, castration-sensitive prostate cancer (CSPC) and castration-resistant prostate cancer (CRPC) cells bearing the following major gene regulatory networks (GRNs) — "TCF7L2+ WNT", "MAFG+", "IRF2+ Inflammatory", "SOX2/4+ Embryonic EMT", "FOSL1+ AP-1", "AR+ HOXB13+", "AR+ HOXB13−", "AR+ IRF+ Inflammatory", "AR+ GI", and "AR+ HOXB13+ FOXA1+" — were classified as PRAD. NEPC cells bearing GRNs "NEPC-A" or "NEPC-A/SOX6" were classified as NEPC-A, and those bearing "NEPC-N" were classified as NEPC-N. Average expression of *MYC*, *MYCN*, and *MYCL* across the three subtypes was visualized using the ’DotPlot’ function in the Seurat package^104^.

### CUT&RUN and ATAC-seq analysis

Bcl files containing raw sequencing data were converted to fastq format, adaptor trimmed, and demultiplexed using bcl2fastq2. Quality of the sequencing data was verified using FastQC. Paired-end sequencing reads were aligned to the mouse genome (GRCm38.p6/mm10) using Bowtie2 (v2.3.4.1)^105^ with ‘‘–local –very-sensitive-local –phred33 –no-unal –no-mixed –no-discordant -I 10 -X 700 -S’’. For CUT&RUN, reads were also aligned to the drosophila genome (dm6) with additional ‘‘–no-overlap’’ to map spike-in reads. Sam files were converted to bam files, sorted, and indexed using samtools (v1.9)^106^. Peaks were called for each individual dataset using MACS2 (v2.2.6)^107^ callpeak function with ‘‘-f BAMPE –keep-dup 1 -q 0.05’’. Intersection of peak sets were performed using the intersectBed program from bedtools (v2.27.1)^108^. Motif analysis was performed using the findMotifsGenome program from HOMER (v4.11)^109^ with ‘‘-size 500 -S 20 -mask’’. Bigwig pileup files were generated using the bamCoverage program from DeepTools (v3.2.1)^110,111^ with ‘‘-bs 10 –extendReads – normalizeUsing RPKM –exactScaling –skipNonCoveredRegions’’ and filtered for mouseENCODE blacklisted genomic regions (v2)^111^ using ‘‘-bl’’ option. For spike-in normalization in perturbation experiments (degradation or knockout), the total number of mapped reads to mouse and *drosophila* genomes was used to generate scale factors, which were applied to the ‘‘–scaleFactor’’ option of bamCoverage for cell number-based normalization. Genome browser tracks of bigwig pileups were generated using IGV (v2.5.1)^112^. Heatmaps and average profiles of bigwig pileups at peaksets were generated using the computeMatrix, plotProfile and plotHeatmap functions from DeepTools. Overlapping analysis of CUT&RUN peaks were generated using Venn module or pairwise intersection module with “--compute frac” of the intervene program (v0.6.5)^113^.

Area-proportional Venn diagrams were generated using BioVenn (https://www.biovenn.nl/index.php)^114^. Peak annotation (association with genes) was performed using GREAT (v4.0.4) (https://great.stanford.edu/great/public/html/)^115^ and gene set over-representation analysis (ORA) was performed using the enricher function from the clusterProfiler R package.

For chromatin state analysis, ChromHMM (v1.23)^116^ was used to build a 9-state chromatin model using CUT&RUN data of 6 histone modifications (H3K4me1, H3K4me3, H3K27ac, H3K36me3, H3K9me3 and H3K27me3) from WT organoids. This model and CUT&RUN the of 6 histone modifications from all 5 organoid lines (WT, AD1, AD2, NE1 and NE2), were used to make chromatin state segmentation of the genome for each line. The chromHMM commands BinarizeBam, LearnModel and MakeSegmentation were all run using default parameters. We picked the 9-state chromatin model since it was the most biologically interpretable model. To quantify the chromatin state changes between organoid lines we used bedtools (v2.27.1) with “intersect -wo” to count the total number of bases that overlap between states and normalized them against the total number of bases in each state in the control line (WT). Heatmaps were created using the ComplexHeatmap R package, and the Sankey plots were created using the ggsankey R package (https://github.com/davidsjoberg/ggsankey).

### Quantification and statistical analysis

Calculations and statistical analyses were performed using Microsoft Excel, GraphPad Prism and R. The statistical details of the experiments are provided in the figure legends and method details wherever applicable.

### Data and code availability

Next generation sequencing data have been deposited at GEO: GSE325159 and will be publicly available as of the date of publication. This study does not report original code. Analyses were performed with published tools as described in the Methods section.

---

**Figure S1.**
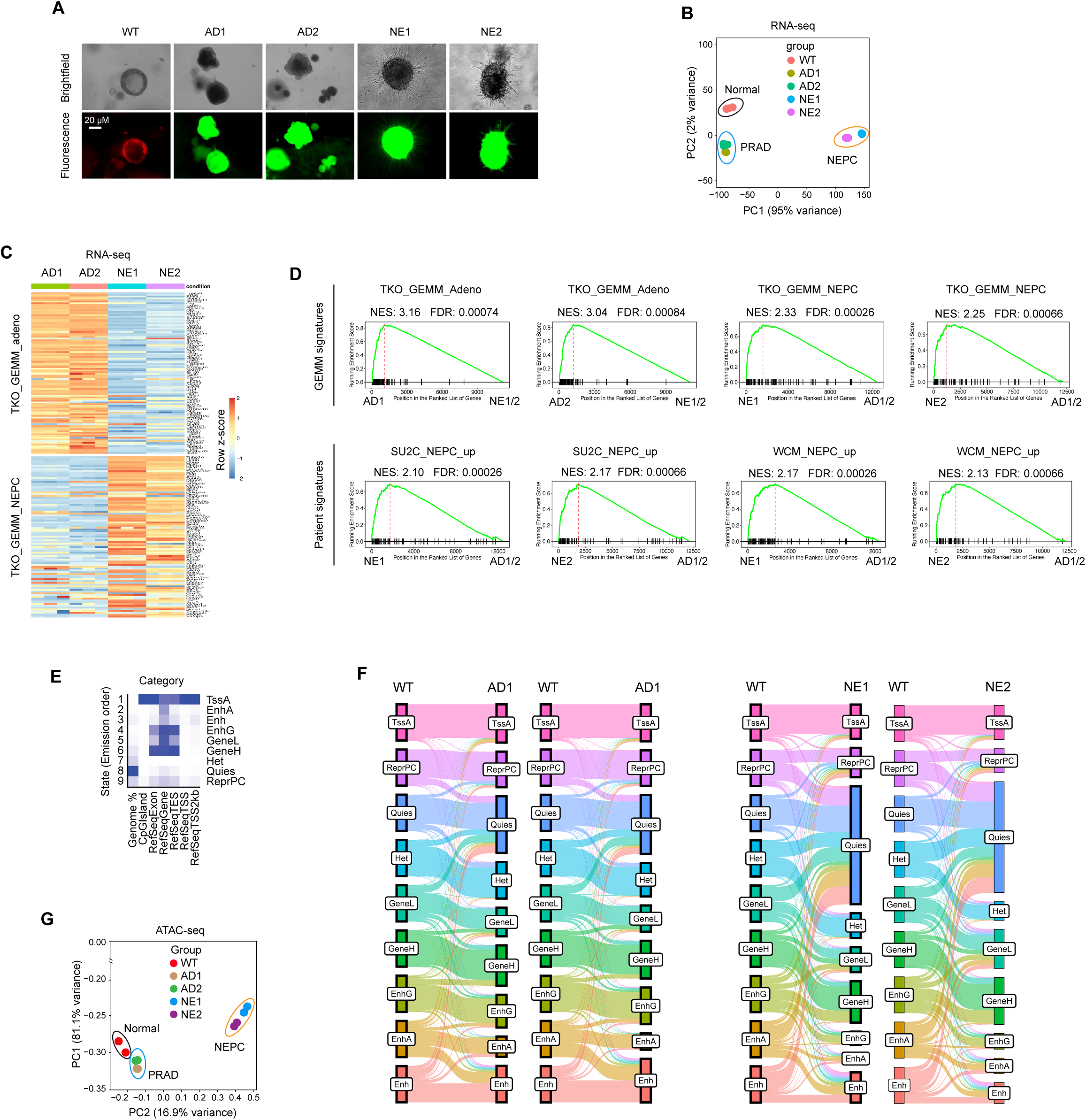
Extensive enhancer reprogramming during PRAD to NEPC transition. Related to Figure 1. (**A**) Representative brightfield and fluorescent microscopy images of indicated organoid lines. Scale bar representing 20 μm applies to all panels. (**B**) Principal component analysis (PCA) of RNA-seq using WT, PRAD and NEPC organoids. (**C**) Heatmap showing z-score transformed expression (RNA-seq) of TKO GEMM PRAD and NEPC signature genes in indicated organoid lines. PRAD and NEPC signatures were the top 100 upregulated genes from scRNA-seq of TKO GEMM tumors in PRAD or NEPC cells compared to all cells of other lineages/states. (**D**) GSEA enrichment plots of TKO GEMM (top) and patient (bottom) PRAD and NEPC signatures in indicated PRAD and NEPC organoid lines. Related to Figure 1E. NES, normalized enrichment score. (**E**) Heatmaps showing the overlap enrichment of indicated chromatin states (Figure 1F) in genomic categories, as calculated by chromHMM. (**F**) Quantification of chromatin state changes comparing AD1, AD2, NE1, NE2 against WT as control. Sankey plots showing the proportion of each chromatin state (chromoHMM) in each organoid line as shown by the height of the rectangles, and the proportion of overlap (connecting ribbons) between the states in the organoid lines compared. (**G**) PCA of ATAC-seq at all peaks in WT, PRAD and NEPC organoids.

**Figure S2.**
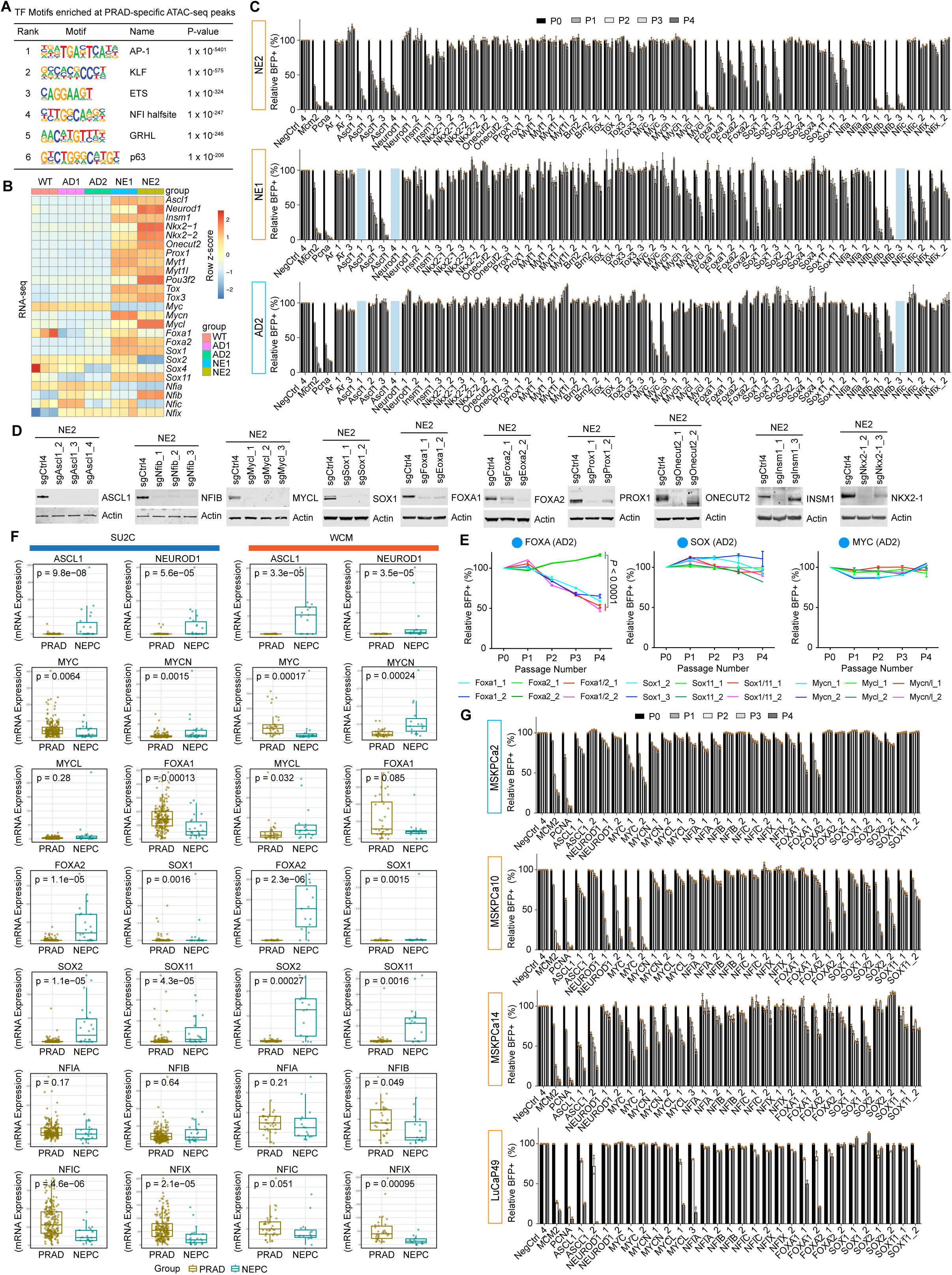
Network of transcription factor dependencies in murine and human NEPC. Related to Figure 2. (**A**) Top 5 *de novo* TF motifs enriched at PRAD-specific ATAC-seq peaks (target coverage > 10%, ranked by p values). (**B**) Heatmap showing z-score transformed expression (RNA-seq) of candidate TFs for CRISPR screen in indicated organoid lines. (**C**) Cell competition assays using at least two independent sgRNAs targeting the DNA binding domains of 25 TF candidates in indicated organoid lines, showing relative BFP percentage over 4 passages (P0-4) after sgRNA transduction. Related to Figure 2D. BFP percentages were normalized to P0 (day 2) and the NegCtrl4 sgRNA at the respective day (relative %BFP+). Data represent mean ± SD from n = 2 technical replicates. (**D**) Western blots of whole cell extract from NE2 organoids with knockout of indicated TFs. Actin was used as loading control. (**E**) Cell competition assays using two independent single or dual sgRNA vectors targeting either or both TF paralogs in indicated PRAD tumoroids, showing relative BFP percentage over 4 passages (P0-4). Related to Figure 2E. BFP percentages were normalized to day 2 (P0) and the NegCtrl4 sgRNA at the respective day (relative %BFP+). Data represent mean ± SD from n = 2 technical replicates. *P* values were calculated using two-tailed unpaired Student’s *t*-tests. (**F**) Boxplots showing expression (bulk RNA-seq) of indicated TFs in tumor samples of the SU2C and WCM prostate cancer patient cohorts. *P* values were calculated using two-tailed Wilcoxon rank sum test. (**G**) Cell competition assays using at least two independent sgRNAs targeting the DNA binding domains of indicated TFs in indicated PDO lines, showing relative BFP percentage over 4 passages (P0-4) after sgRNA transduction. Related to Figure 2H. BFP percentage was measured only on P0, P2 and P4 for LuCaP49. BFP percentages were normalized to P0 (day 2) and the NegCtrl4 sgRNA at the respective day (relative %BFP+). Data represent mean ± SD from n = 2 technical replicates.

**Figure S3.**
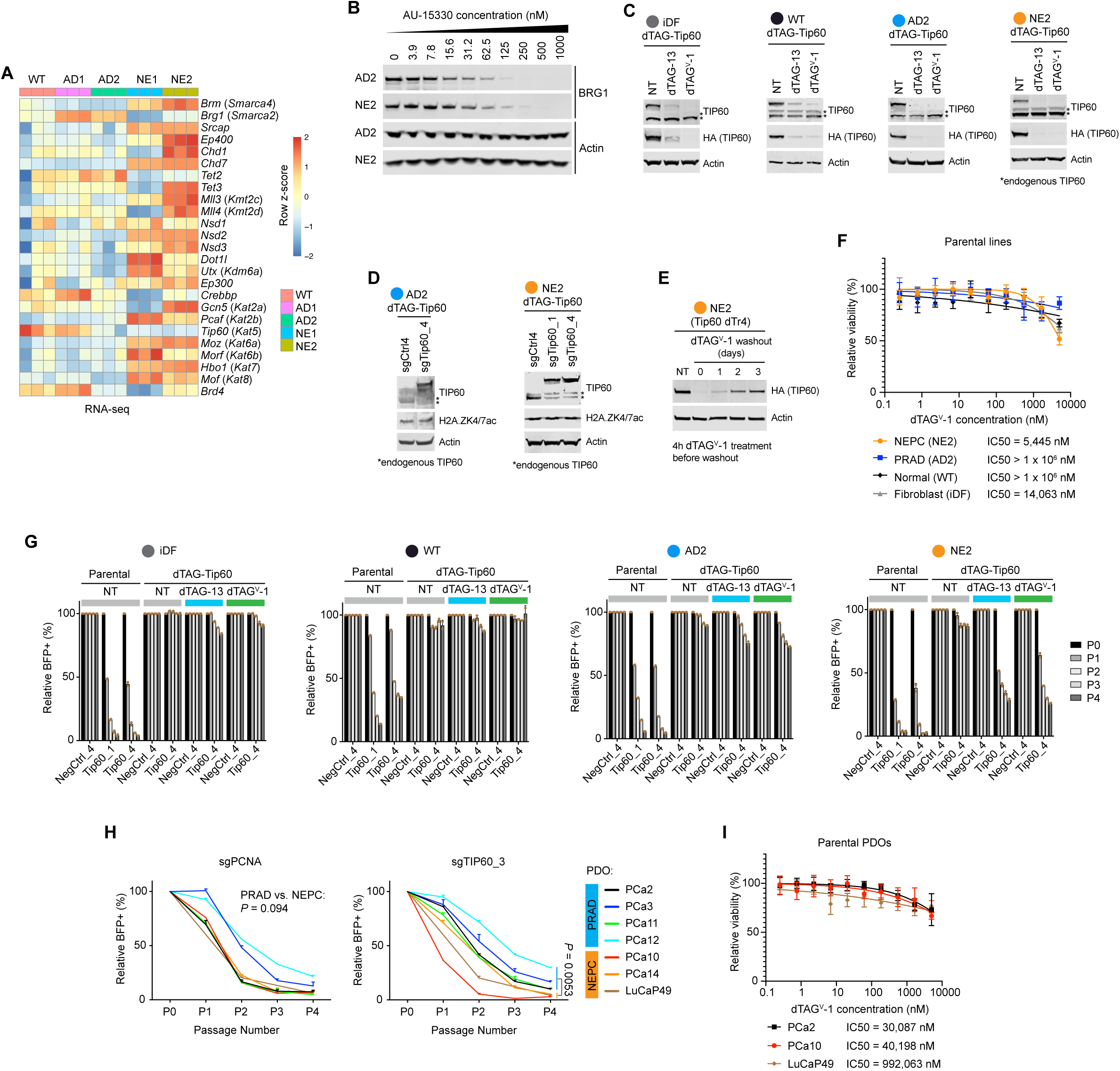
Enhanced dependency on the TIP60 coactivator complex in NEPC. Related to Figure 3. (**A**) Heatmap showing z-score transformed expression (RNA-seq) of candidate chromatin coactivators for CRISPR screen in indicated organoid lines. (**B**) Western blots of whole cell extract from AD2 and NE2 organoids treated with a 2-fold gradient of AU-15330 for 24 hours. In (**B**), (**C**), (**D**) and (**E**), Actin was used as loading control. (**C**) Western blots of whole cell extract of indicated cell/organoid lines stably overexpressing dTAG-Tip60 and treated with dTAG-13 or dTAG^V^-1 for 24 h. (**D**) Western blots of whole cell extract of AD2 and NE2 organoids stably overexpressing dTAG-Tip60 and with knockout of endogenous *Tip60* using indicated sgRNAs. (**E**) Western blots of whole cell extract of NE2 *Tip60* dTr4 organoids pretreated with dTAG^V^-1 for 4 h and then washed out of dTAG^V^-1 for indicated days. For washout, organoids were extracted from the Matrigel and reseeded in organoid culture conditions without dTAG^V^-1. (**F**) dTAG^V^-1 dose response curves and IC50 values for indicated parental lines (no dTAG-*Tip60* or *Tip60* sgRNA). PDOs were treated with the same dTAG^V^-1 gradient as in Figure 3E for 6 days before CellTiter Glo assays were performed to determine cell viability. Data represent mean ± SD from n = 10 biological replicates. (**G**) Cell competition assays using two independent *Tip60* sgRNAs in indicated cell/organoid lines stably overexpressing dTAG-Tip60 and treated with dTAG-13 or dTAG^V^-1 for the duration of the experiment, showing relative BFP percentage over 4 passages (P0-4). BFP percentages were normalized to day 2 (P0) and the NegCtrl_4 sgRNA at the respective day (relative %BFP+). Data represent mean ± SD from n = 2 technical replicates. NT, non-treated. (**H**) Cell competition assays using two independent *TIP60* sgRNAs in indicated PRAD and NEPC PDOs, showing relative BFP percentage over 4 passages (P0-4). Related to Figure 3H. BFP percentage was measured only on P0, P2 and P4 for LuCaP49. BFP percentages were normalized to day 3 (P0) and the NegCtrl_4 sgRNA at the respective day (relative %BFP+). Data represent mean ± SD from n = 2 technical replicates. *P* values were calculated using two-tailed unpaired Student’s *t*-tests. (**I**) dTAG^V^-1 dose response curves and IC50 values for indicated parental PDO lines (no dTAG-*Tip60* or *TIP60* sgRNA). PDOs were treated with the same dTAG^V^-1 gradient as in Figure 3J for 10 - 13 days before CellTiter Glo assays were performed to determine cell viability. Data represent mean ± SD from n = 10 biological replicates.

**Figure S4.**
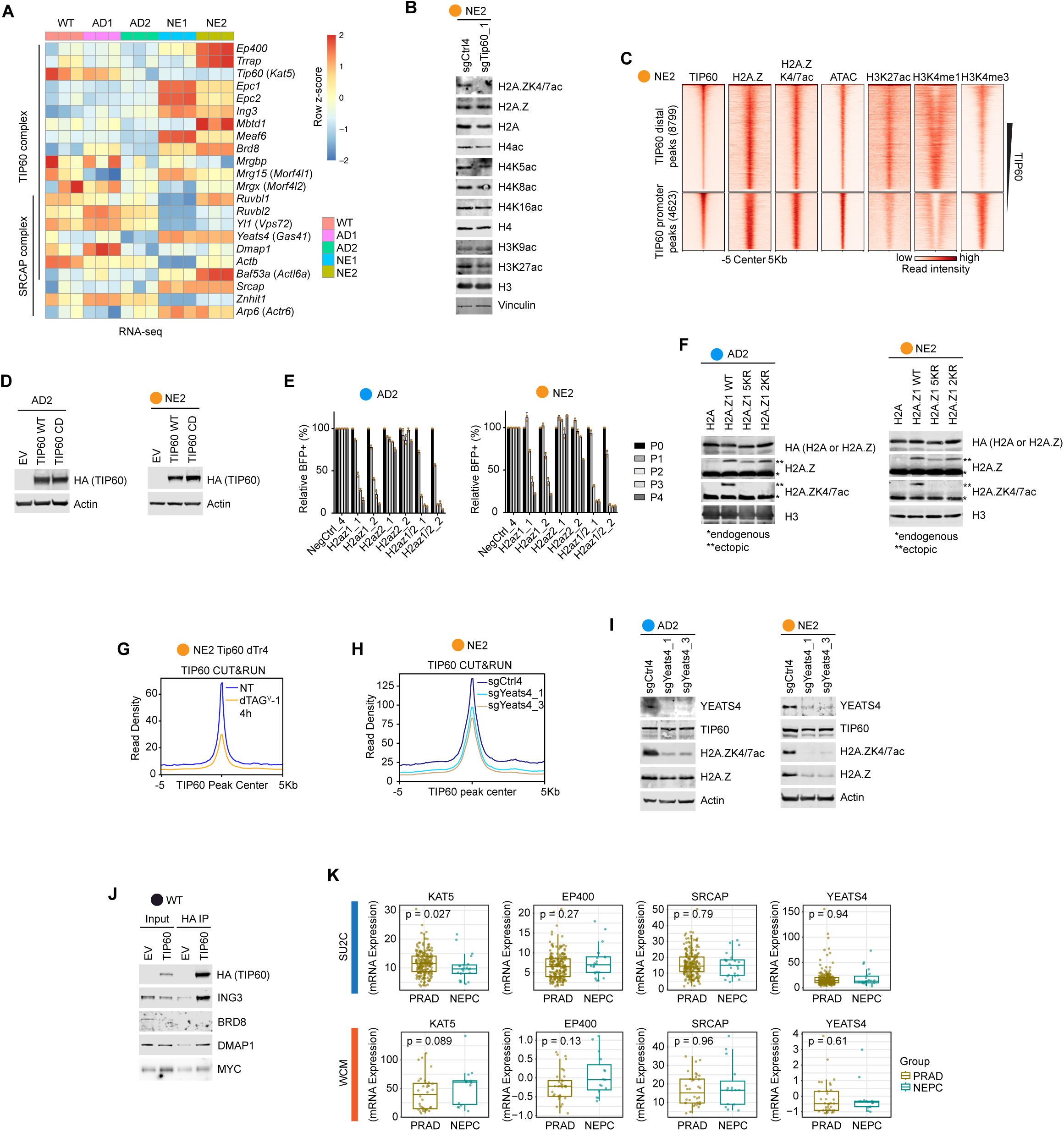
TIP60-C dependency in NEPC is mediated through H2A.Z acetylation. Related to Figure 4. (**A**) Heatmap showing z-score transformed expression (RNA-seq) of subunits of the TIP60 and SRCAP complexes in indicated organoid lines. (**B**) Western blots of whole cell extract from NE2 organoids with *Tip60* knockout showing levels of H4 and H2A.Z acetylation. H3 acetylation were used as negative controls. Vinculin was used as loading control. (**C**) Heatmap showing ATAC-seq and occupancy (CUT&RUN) of indicated proteins and histone modifications at all TIP60 peaks (promoter and distal) in NE2 organoids. (**D**) Western blots of whole cell extract from AD2 and NE2 organoids stably overexpressing HA-tagged wild type (WT) or Q377E/G380E catalytically dead (CD) TIP60. Actin was used as loading control. (**E**) Cell competition assays using two independent single or dual sgRNA vectors targeting either or both H2A.Z isoforms in AD2 and NE2 organoids. Data represent relative BFP percentage over 4 passages (P0-4). BFP percentages were normalized to day 2 (P0) and the NegCtrl_4 sgRNA at the respective day (relative %BFP+). Data represent mean ± SD from n = 2 technical replicates. (**F**) Western blots of whole cell extract from AD2 and NE2 organoids expressing HA-tagged canonical H2A or WT/mutant H2A.Z1. 2KR, K4/7R mutant. 5KR, K4/7/11/13/15R mutant. H3 was used as loading control. (**G**) Average profile showing occupancy (CUT&RUN) of TIP60 (via anti-TIP60 antibody) at TIP60 peaks in NE2 dTAG-Tip60 (dTr4) organoids treated with 500 nM dTAG^V^-1 for 4 h. NT, non-treated. (**H**) Average profile showing occupancy (CUT&RUN) of TIP60 in NE2 organoids with *Yeats4* knockout. NT, non-treated. (**I**) Western blots of whole cell extract from AD2 and NE2 organoids with *Yeats4* knockout. (**J**) Western blot analyses of immunoprecipitation with HA tag in WT organoids expressing HA-FLAG-tagged TIP60 (Tip60 dTr4) or empty vector (EV). (**K**) Boxplots showing expression (bulk RNA-seq) of indicated subunits of the TIP60 and SRCAP complexes in tumor samples of the SU2C and WCM prostate cancer patient cohorts. *P* values were calculated using two-tailed Wilcoxon rank sum test.

**Figure S5.**
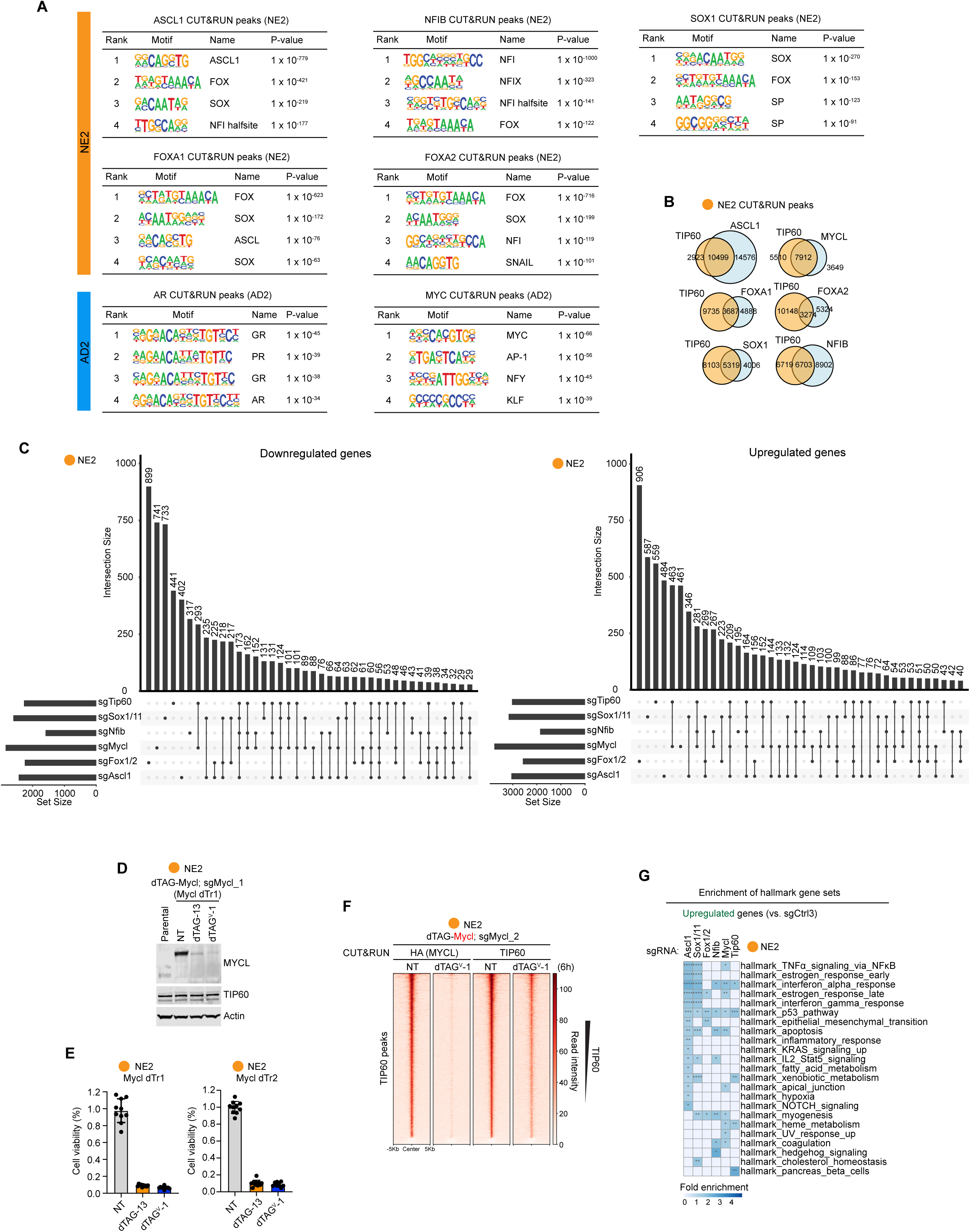
NEPC TFs function as a network, culminating in chromatin recruitment of TIP60-C by MYCL. Related to Figure 5. (**A**) Top 4 *de novo* TF motifs enriched at CUT&RUN peaks of indicated TFs in AD2 or NE2 organoids (target coverage > 10%, ranked by p values). (**B**) Venn diagram depicting overlap of TIP60 CUT&RUN peaks with peaks of indicated TFs. Related to Figure 5B. (**C**) UpSet plot showing intersections of significantly downregulated (FDR < 0.05, log2FC < 0) and upregulated (FDR < 0.05, log2FC > 0) genes upon 3-day knockout of indicated factors in NE2 organoids. (**D**) Western blots of NE2 organoids with expression of dTAG-Mycl and knockout of endogenous *Mycl* (sgMycl_1) alleles. Organoids were treated with dTAG-13 or dTAG^V^-1 for 24 h. Actin was used as loading control. (**E**) Cell viability of NE2 organoids with expression of dTAG-Mycl and knockout of endogenous *Mycl* alleles using sgMycl_1 (dTr1) or sgMycl_2 (dTr2) after 6 days of treatment of 500 nM dTAG-13 or dTAG^V^-1. Data represent mean ± SD from n = 10 biological replicates. (**F**) Heatmap showing occupancy (CUT&RUN) of MYCL (via HA tag) and TIP60 at TIP60 peaks in NE2 organoids with expression of dTAG-Mycl and knockout of endogenous *Mycl* alleles. Related to Figure 5E. Organoids were treated with 500 nM dTAG^V^-1 for 6 h. NT, non-treated. (**G**) Overrepresentation analyses (FDR < 0.05) of hallmark gene sets in upregulated genes (FDR < 0.05, log2FC < 0) upon 3-day knockout of indicated genes compared to control sgRNA in NE2 organoids. Related to Figure 5H. ****, FDR < 0.0001. ***, FDR < 0.001. **, FDR < 0.01. *, FDR < 0.05.

**Figure S6.**
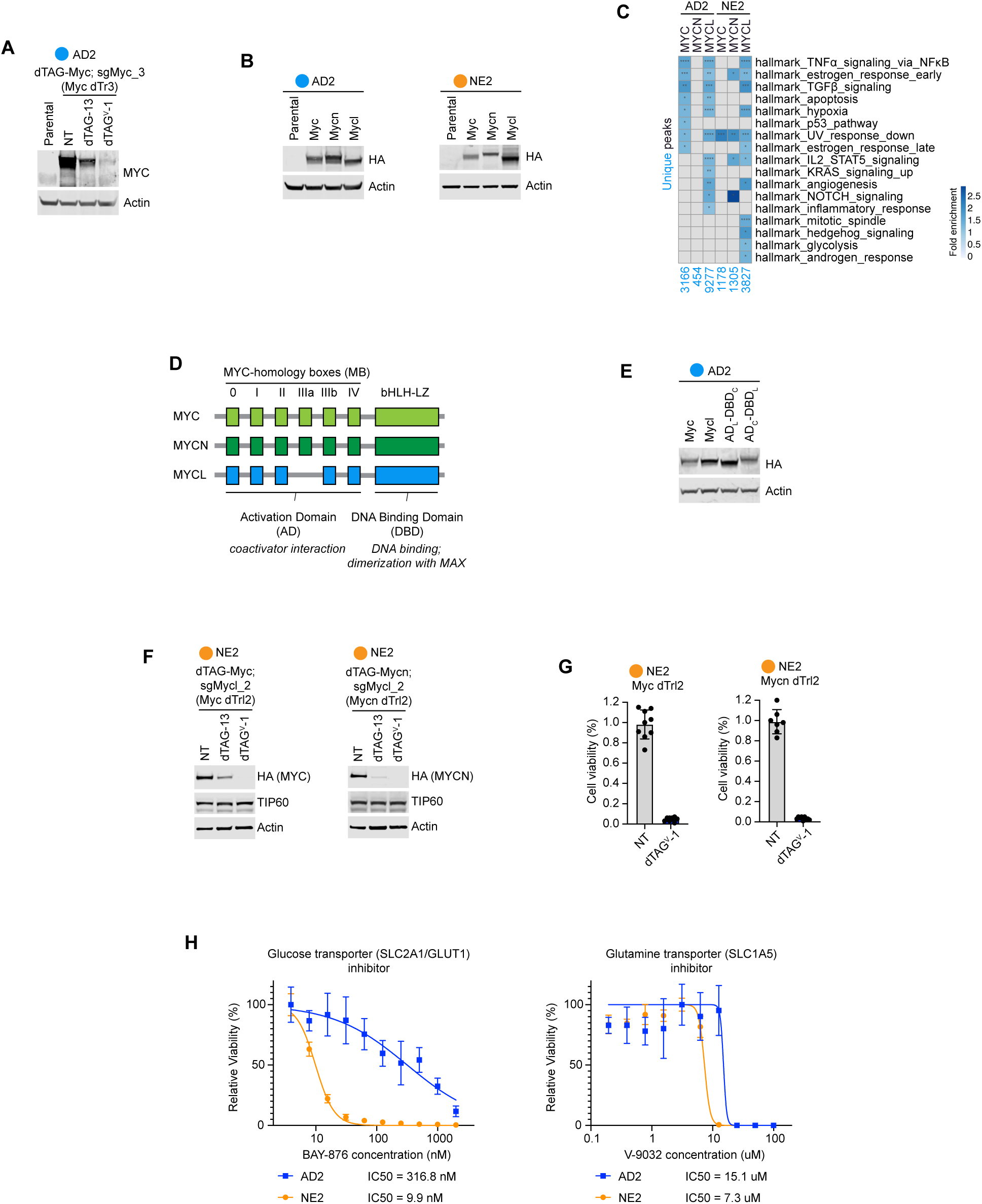
MYC paralogs partner with TIP60 for coactivation exclusively in the NEPC lineage of prostate cancer. Related to Figure 6. (**A**) Western blots of AD2 organoids with expression of dTAG-Myc and knockout of endogenous *Myc* (sgMyc_3) alleles. Organoids were treated with dTAG-13 or dTAG^V^-1 for 24 h. In (**A**), (**B**), (**E**) and (**F**), Actin was used as loading control. (**B**) Western blots of whole cell extract from AD2 and NE2 organoids overexpressing indicated *Myc*, *Mycn* or *Mycl* (with HA**-**FLAG-dTAG). (**C**) Overrepresentation analysis (FDR < 0.05) of hallmark gene sets (50 sets) in genes associated with unique CUT&RUN peaks of MYC, MYCN and MYCL in AD2 and NE2 organoids. Related to Figure 6F. ****, FDR < 0.0001. ***, FDR < 0.001. **, FDR < 0.01. *, FDR < 0.05. (**D**) Schematic showing the domain structures and functions of the three MYC paralogs. (**E**) Western blots of whole cell extract from AD2 organoids overexpressing wild type and chimeric MYC proteins (with HA**-**FLAG-dTAG). (**F**) Western blots of whole cell extract from NE2 organoids with expression of dTAG-Myc or dTAG-Mycn and knockout of endogenous *Mycl* alleles using sgMycl_2. Organoids were treated with 500 nM dTAG-13 or dTAG^V^-1 for 24 h. (**G**) Cell viability of NE2 *Myc* dTrl2 (dTAG-Myc; sgMycl_2) and *Mycn* dTrl2 (dTAG-Mycn; sgMycl_2) organoids after 6 days of treatment of 500 nM dTAG^V^-1. Data represent mean ± SD from n = 9 (*Myc* dTrl2) and n = 7 (*Mycn* dTrl2) biological replicates. (**H**) BAY-875 (glucose transporter inhibitor) and V-9032 (glutamine transporter inhibitor) dose response curves and IC50 values for NEPC (NE2) vs. PRAD (AD2). Tumoroid lines were treated with the indicated drug gradients for 6 days before CellTiter Glo assays were performed to determine cell viability. Data represent mean ± SD from n = 10 biological replicates.

